# Insect olfaction-inspired biohybrid sensor array for selective airborne pheromone detection and early monitoring of invasive *Rhynchophorus ferrugineus*

**DOI:** 10.64898/2026.08.13.744605

**Authors:** Khasim Cali, Binu Antony, Corrado Di Natale, Alexandro Catini, Nicolas Montagné, Emmanuelle Jacquin-Joly, Mohammed A. AlSaleh, Yousef Al-Fehaid, Krishna C. Persaud, Arnab Pain

## Abstract

The red palm weevil, *Rhynchophorus ferrugineus* (Olivier) (Coleoptera: Curculionidae), is a globally invasive quarantine pest threatening palm cultivation across 49 countries and inflicting annual economic losses estimated at over USD 100 million. Weevil larvae burrow into palm trunks, causing progressive internal structural damage that rarely produces visible external symptoms until lethal injury has occurred, rendering early detection exceptionally challenging. In the absence of effective early-warning surveillance technologies, tens of thousands of infested palm trees have been removed across major palm-cultivation regions in the Middle East and Mediterranean basin. Rapid, sensitive detection of volatile organic compounds (VOCs) emitted by weevil colonies and infested palm trees therefore represents a critical unmet need for timely pest surveillance and intervention. Existing artificial gas sensors lack the chemical selectivity required to discriminate among structurally similar VOCs, and no validated field-deployable early-detection platform has been established to date. Here, we report a portable biohybrid sensor array that mimics insect olfaction by exploiting two classes of diagnostic chemical signatures: the male-released aggregation pheromone (4RS,5RS)-4-methylnonan-5-ol (ferrugineol) and ethyl ester volatile blends emitted by weevil-infested palm trees. The *R. ferrugineus* odorant receptor RferOR1 was stabilised in lipid nanodiscs and co-immobilised with two *in vivo*-synthesised odorant-binding proteins (RferOBP1768 and RferOBP23) on quartz crystal microbalance (QCM) transducers to construct the biohybrid sensing platform. The sensor array achieved selective detection of airborne ferrugineol at a limit of detection of approximately 60 parts per billion (ppb) under field conditions, distinguishing infested from healthy palms. OBP- and OR-functionalised sensors retained full functional activity for 12 and 7 months, respectively, under ambient storage, confirming operational robustness and shelf life suitable for long-term field deployment. This work translates the molecular architecture of the insect olfactory system into a practical, field-validated chemical sensor platform with direct applicability to early-stage *R. ferrugineus* infestation monitoring and sustainable integrated pest management.

**Graphical abstract:** 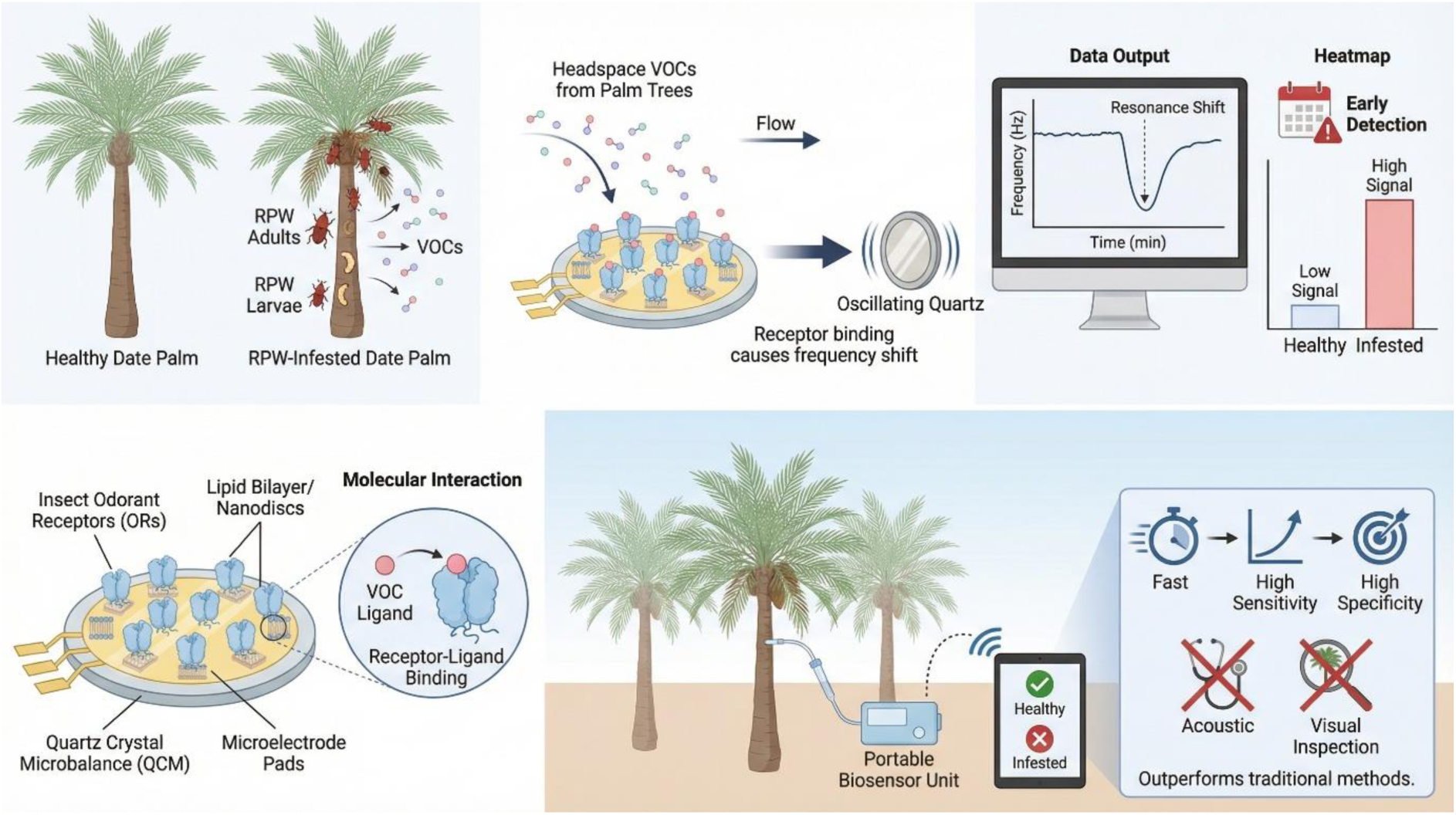

## 1. Introduction

The accelerating pace of globalisation over the past five decades has facilitated the introduction and establishment of numerous invasive insect species in novel geographic ranges, where they have become destructive agricultural pests causing severe ecological and economic damage worldwide [1]. A direct consequence of the global palm trade has been the rapid spread of the red palm weevil (RPW), *Rhynchophorus ferrugineus* (Coleoptera: Curculionidae), and the South American palm weevil, *R. palmarum* [2]. Both species are listed as quarantine pests of A2 risk status by the Food and Agriculture Organisation (FAO) of the United Nations and the European and Mediterranean Plant Protection Organisation (EPPO), designating them as established but incompletely distributed pests requiring strict regulatory control to prevent further spread. Early detection of palm weevil infestation remains elusive with existing surveillance tools. Here, exploiting biorecognition elements derived from the weevil’s chemosensory system, we demonstrate a biohybrid biosensor capable of sensitively detecting volatile aggregation pheromones associated with infested palm trees.

*R. ferrugineus* is native to South and Southeast Asia and Melanesia, from where it has spread across the Arabian Peninsula, South Asia, and the Mediterranean, now threatening palm ecosystems — including UNESCO World Heritage Sites such as Socotra Island and iconic cultural landscapes such as the French Riviera — across 49 countries. Over the past 30 years, its host range has expanded from four palm species in the 1950s to over 40 by 2020 [2], encompassing date, coconut, oil, Canary Island, and Washingtonia palms. The weevil currently threatens palm cultivation across the Near East, North Africa, the Gulf states, much of Europe, 28 Asian countries, Central America, and the Caribbean [2], including iconic palms in UNESCO World Heritage sites (e.g., Socotra Island) and cultural landscapes (e.g., the French Riviera). *R. palmarum* is an equally destructive pest of commercial (coconut and oil palm) and ornamental (Canary Island date palm) palms across its native range in Central and South America and parts of Mexico, as well as invaded regions including California, USA. Critically, *R. palmarum* serves as the biological vector of the red ring nematode, *Bursaphelenchus cocophilus*, which causes red ring disease — a lethal, incurable palm disease [2]. The recent invasion of R. palmarum into southern California has resulted in the death of over 20,000 urban palms [2, 3] and threatens to spread into the economically vital date palm production areas of the Coachella Valley. The establishment of *R. palmarum* in the USA parallels the invasion trajectory of *R. ferrugineus*, underscoring the urgent need for effective early-detection and monitoring technologies applicable to both species.

Once weevils infest a palm, larvae remain concealed within the trunk, feeding destructively on internal tissues. Without early detection, infestations are invariably fatal to the host tree. Currently available detection methods — including visual inspection, pheromone-baited trapping, and trained detection dogs — are labour-intensive, costly, and unreliable for early-stage detection [2]. Emerging approaches such as optical sensors [4], acoustic monitoring [5], and machine learning-based classification systems [6] remain constrained by limitations in accuracy, cost, and field deployability [7]. Detection of airborne pheromones from infested palms has been demonstrated under experimental conditions, offering a promising avenue for non-invasive early surveillance [8]; however, odour-based sensor platforms for field deployment have not yet been realised [9].

The ecological success and invasive capacity of *R. ferrugineus* is substantially attributable to its sophisticated chemical communication system. Male RPWs release aggregation pheromones — (4RS,5RS)-4-methylnonan-5-ol (ferrugineol) and (4RS,5RS)-4-methylnonan-5-one (ferrugineone) — that attract individuals of both sexes [10]. *R. palmarum* males produce a structurally distinct aggregation pheromone, (4S,2E)-6-methylhept-2-en-4-ol (rhynchophorol) [11]. For both species, synthetic blends of the (S)- and (R)-enantiomers of these pheromone components are widely employed in pheromone-baited traps, and their attractiveness is substantially enhanced by the addition of host plant material such as decaying palm stem tissue, reflecting synergistic interactions between pheromone and host plant volatiles [12]. Notably, both species show olfactory responses to ferrugineol and ferrugineone. The odorant receptor mediating detection of these pheromone components has been characterised in *R. ferrugineus* as RferOR1 [13] and its orthologue RpalOR1 identified in *R. palmarum* [14]. These pheromone signals, in combination with palm tissue volatiles and fermentation-derived kairomones from damaged tissue, coordinate weevil attraction to infested trees. Electroantennogram studies have further demonstrated that both species’ antennae respond to a range of palm-associated volatiles, including short-chain ethyl esters — ethyl acetate, ethyl propionate, and ethyl butyrate—collectively termed palm esters [12], which facilitate coordinated mass attack behaviour.

Insects detect environmental volatiles through two principal classes of chemosensory proteins: odorant-binding proteins (OBPs) and odorant receptors (ORs). OBPs are soluble, extracellular proteins that bind and transport hydrophobic odorant molecules through the aqueous sensillar lymph to the receptor membrane, while ORs are membrane-bound ligand-gated ion channels that transduce odorant binding into neuronal signals [13, 15, 16]. The high physicochemical stability of OBPs has made them attractive candidates for immobilisation on transducer surfaces for volatile compound detection [17, 18]. Importantly, OBPs immobilised via self-assembled monolayers at the gate of organic bioelectronic transistors have been shown to enable sensitive, quantitative discrimination of weak interactions with neutral enantiomers that bind differentially to the protein [19]. ORs form obligate heteromeric complexes with a conserved co-receptor (Orco) and are characterised by high ligand specificity and structural diversity [20]. OR-based biosensors have been developed using Drosophila receptors [21], and OBP-based sensors derived from honeybee proteins have demonstrated detection of floral and pheromone volatiles *via* electrochemical impedance spectroscopy [22, 23]. Despite long-standing interest in insect antenna-based biosensing, practical deployment has been impeded by challenges in OBP–OR array integration and incomplete functional characterisation of insect ORs [24].

The majority of existing chemical biosensors operate in the liquid phase. In the present study, we developed a novel biohybrid biosensor for the detection of airborne volatiles using quartz crystal microbalances (QCMs), piezoelectric transducers that measure nanogram-scale mass changes arising from molecular binding events [25]. We designed a sensitive, biomimetic sensor array that recapitulates key functional properties of the insect antenna by integrating both OBPs and ORs as biorecognition elements. This was made possible by the prior identification and functional characterisation of *R. ferrugineus* chemosensory proteins, including the pheromone receptor RferOR1, the plant volatile receptor RferOR2, the obligate co-receptor RferOrco, and multiple odorant-binding proteins (RferOBPs) [12, 13, 15, 16, 26]. Our strategy was to develop and express variants of these proteins with differing binding affinities for target volatile compounds, creating a sensor array capable of discriminating pheromone volatiles from background plant volatiles and kairomones. The functionalised sensor array selectively and sensitively detects airborne ferrugineol and ferrugineone vapours emanating from weevil-infested palms by mimicking the native functions of RferOR1 and RferOBPs. To address the inherent instability of membrane-bound ORs in biosensor applications, we embedded RferOR1 within lipid nanodiscs, which stabilised the receptor in a native-like conformation and preserved ligand-binding activity. We further evaluated sensor array shelf life under ambient storage conditions and validated field applicability in palm plantation environments. This portable, biomimetic detection platform offers a practical and sustainable tool for early detection and real-time monitoring of *R. ferrugineus* infestation, with direct relevance to integrated pest management programmes.

## 2. Methods

### 2.1 Chemicals and pheromones

Synthetic palm volatiles (Table 1) and pheromone were obtained from Sigma-Aldrich (Saint-Louis, MO, USA). Beetle pheromones and all related compounds were in our lab stock and were prepared as previously described [13, 14].

**Table 1.** Volatile compounds used to characterise the biosensor array, with chemical purity, CAS registry number, abbreviation, saturated vapour pressure, and biological source of emission [12–14].

| Chemical | Purity | CAS | Abbreviation | Vapor pressure [Pa] | Source of emission |
| --- | --- | --- | --- | --- | --- |
| (4RS,5RS)-4-methylnonan-5-ol | 99 % | 154170-44-2 | Ferrugineol | 7.4 | Red palm weevil |
| 4(RS)-methylnonan-5-one | 98 % | 35900-26-6 | Ferrugineone | 45 | Red palm weevil |
| nonan-5-ol | 99 % | 623-93-8 | nonan-5-ol | 1729 | Red palm weevil |
| nonan-5-one | 97 % | 502-56-7 | nonan-5-one | 73 | Red palm weevil |
| 5-methyloctan-4-one | 98 % | 6175-51-5 | Cruentol | 1333 | Palmetto weevil (*R. cruentatus*) |
| ethyl hexanoate | >99% | 123-66-0 | Ethyl hexanoate | 207 | *E. guineensis* fermented sap; *C. nucifera* crown; *P. canariensis* affected stem and fermented leaves |
| ethyl valerate | 99% | 539-82-2 | Ethyl valerate | 299 | *P. canariensis* affected stem |
| butyl butyrate | 98% | 109-21-7 | Butyl butyrate | 1378 | *P. canariensis* affected stem |
| propyl butyrate | >95% | 105-66-8 | Propyl butyrate | 793 | *P. canariensis* affected stem |
| ethyl butyrate | 99% | 105-54-4 | Ethyl butyrate | 1999 | *C. nucifera* crown, *E. guineensis* fermented sap, trunk; *P. canariensis* affected stem; *S. officinarum* stalk |
| ethyl propionate | 99% | 105-37-3 | Ethyl propionate | 199 | *E. guineensis* trunk; *P. canariensis* affected stem, fermented leaves; *S. officinarum* stalk |
| ethyl acetate | 99.8% | 141-78-6 | Ethyl acetate | 9732 | *P. canariensis* healthy and affected stem, fermented leaves; *P. dactylifera* fruits |
| 2(E)-6-methyl-2-hepten-4-ol | 99% | 1569-60-4 | Rhynchophorol | 31.46 | American palm weevil (*R. palmarum*) |

### 2.2 Red palm weevil rearing

RPWs used in this study came from lab cultures and field collections. Lab-reared RPWs were maintained on sugarcane stems [13] and considered pure lines, unblended with other populations.

### 2.3 Cloning of RferOBP1768, RferOBP23, and RferOR1

We cloned RferOR1 [13] and OBPs RferOBP1768 and RferOBP23 [15] for biosensor development by the following method. Total RNA was extracted from the antennae of 10-day-old RPW adults using the PureLink RNA Mini Kit (Thermo Fisher, USA). First-strand cDNA was synthesized using SuperScript IV Reverse Transcriptase (Thermo Fisher) from 1 µg of total RNA extracted (PureLink RNA Mini Kit, Thermo Fisher). The full-length OBPs and RferOR1 were PCR-amplified and gel-purified using a Wizard SV Gel and PCR Clean-up system (Promega), as described previously [13, 15]. The purified DNA was ligated into a pGEM®-T easy vector and transformed into JM109 *E. coli* cells (Promega). Plasmid DNA was purified using a QIAprep Miniprep kit (Qiagen, Venlo, Netherlands), and the insert was sequenced on an ABI 3500 (Thermos). The plasmids were isolated manually and sequenced in both directions using an ABI 3500 (Thermo Fisher) for sequence verification.

### 2.4 In silico mutagenesis and docking

*In silico* mutagenesis and docking screening were carried out as previously described [17] as follows. Amino acid sequences of RferOBP1768 and RferOBP23 were modelled into 3-D structures using homology modelling tools from the I-TASSER server for protein structure and function prediction. These structures were used as a template for *in silico* mutagenesis experiments. The Computed Atlas of Surface Topography of proteins (CASTp) web server was used to identify potential binding pockets in the proteins. This was followed by single amino acid replacement to identify stable mutations around the ligand binding pocket using a program called PoPMuSiC (Prediction of Protein Mutant Structural Changes). The identified stable mutants were then constructed and visualised using molecular graphics software PyMOL (Schrödinger).

CASTp provides an online resource for locating, delineating and measuring concave surface regions on the three-dimensional structures of proteins. These include pockets located on protein surfaces and voids buried in the interior of proteins. The measurement includes the area and volume of a pocket or void, calculated analytically using a solvent-accessible surface model (Richards’ surface) and a molecular surface model (Connolly surface). CASTp includes a graphical user interface, flexible interactive visualization, and on-the-fly calculations for user-uploaded structures accessed (http://cast.engr.uic.edu9) and as described earlier [17].

The PoPMuSiC software is a tool for the computer-aided design of mutant proteins with controlled stability properties. It evaluates the changes in stability of a given protein or peptide under single-site mutations, based on the protein’s structure. Three modes are available: Systematic, Manual, or File. The Systematic tool evaluates the stability changes resulting from all possible mutations and returns a report containing a list of the most stabilizing or destabilizing mutations, or of the mutations that do not affect stability. The Manual tool predicts the stability change resulting from one or more given mutations. The File tool predicts the stability changes resulting from a list of mutations specified by the user in an uploaded file. For this work, the systematic tool was used to evaluate the stability changes resulting from all possible mutations of RrefOBP1768 and RrefOBPP23 (where each residue was mutated with each of the 20 amino acids minus its own). From the resulting file, we concentrated on the mutations of the main binding pocket residues identified by the CASTp server tool above. All mutations that were identified as stabilizing were selected, recorded, and used in the docking screening process below. A stable mutant must have a negative delta delta energy value (-ΔΔG).

### 2.5 Docking screening

The docking screening was carried out as described [17]. Briefly, the Swissdock server provided by the Swiss Institute of Bioinformatics (http://swissdock.vital-it.ch/docking) using “EADock DSS software was used for the docking of the ligands into the generated mutants and WT. The resulting docking predictions were viewed and analyzed using the Swissdock server plugin in UCSF Chimera. Selected mutants were designed as 6-His tag constructs, and the gene synthesis and protein expression were ordered from GenScript Biotech Corporation (Piscataway, NJ, USA).

### 2.6 Peptide synthesis and binding affinity assays

Wild-type and mutant odorant-binding proteins (OBPs), specifically RferOBP1768_Q12V and RferOBP23_R49L, were cloned, expressed, and purified to support biosensor development. The OBP peptide synthesis was outsourced to GenScript (USA). Briefly, the OBP gene sequences were inserted into the pET-30a (+) expression vector, which carries an N-terminal His tag for subsequent protein purification. These recombinant plasmids were then transformed into the *E. coli* strain BL21 Star™ (DE3). For expression, a single transformed colony for each gene construct was inoculated into Luria-Bertani (LB) medium supplemented with the appropriate antibiotics. The cultures were incubated at 37°C with shaking at 200 rpm. The addition of IPTG induced protein expression, and the progression of expression was monitored using SDS-PAGE.

To maximize protein yield, recombinant BL21 Star™ (DE3) cells preserved in glycerol stocks were further inoculated into 5052 auto-induction medium containing the required antibiotics. Cultures were grown at 37°C until the optical density at 600 nm (OD 600) reached approximately 3. The cultures were then maintained at 37°C for an additional 4 hours to allow for protein expression. After cultivation, cells were harvested by centrifugation, and the resulting pellets were resuspended in lysis buffer. Cell disruption was achieved through sonication, and following further centrifugation, the clarified supernatant was collected for protein purification. The target proteins were purified using a one-step nickel affinity chromatography protocol, taking advantage of the His tag. The purified protein was sterilized by filtration through a 0.22 μm membrane filter before being stored in aliquots for subsequent assays. Protein concentration was determined using the Bradford assay. SDS-PAGE and Western blotting confirmed the purity and molecular weight of the purified proteins.

### 2.7 Fluorescence binding assays

The dissociation constants (K_D_) of the OBPs against target analytes were determined in competitive binding measurements as previously reported [17]. Firstly, the K_D_ of the fluorescence probe N-phenylnaphthalene-1-amine (1-NPN) against the protein was determined. To a 1 µM solution of the protein in 50 mM Tris-HCl, pH 7.4, aliquots of 1 mM 1-NPN in methanol were added to achieve final concentrations of 0 – 16 µM. The probe was excited at 295 nm, and emission spectra were recorded between 337 nm -450 nm (OBP protein–NPN complex peak 405-410nm). The interaction between the protein and the probe was monitored by recording the fluorescence intensity increase upon addition of 1-NPN aliquots. The experiments were replicated at least three times. The dissociation constant (K_D_) of the OBP-protein-NPN complex was calculated from the binding curve by non-linear-least-squares fit of the experimental data using the equation y = B_max_ [NPN] / (K_D_ + [NPN]) where [NPN] is the concentration of the free probe, y is the specific binding derived by measuring fluorescence intensity and B_max_ is the maximum amount of complex formed at saturation. The computer program used to fit the data was SigmaPlot 12.3 (Systat Software, Inc., USA). Once the K_D_ of 1-NPN was determined, the K_D_ of the target analytes was measured in competitive binding assays, recording the fluorescence intensity decrease upon addition of aliquots of 1 mM analytes to give final concentrations between 0 – 16 µM to a solution containing 1 µM protein and 2 µM 1-NPN in 50 mM Tris-HCl, pH 7.4. The K_D_ of the competitor analytes was calculated from the corresponding IC_50_ values (concentrations of the competitor analytes giving half of the initial fluorescence intensity value of 1-NPN) using the equation: K_D_ = [IC_50_]/ (1 + [NPN]/ K_NPN_), [NPN] being the free concentration of 1-NPN and K_NPN_ being the dissociation constant of the protein-probe complex determined [17].

Emission fluorescence spectra were recorded using a Perkin-Elmer LS55 or LS50 Luminescence spectrometer instrument at 25 °C in a right-angle configuration, with a 1-cm light path quartz cuvette and 5-nm slits for both excitation and emission.

### 2.8 Cell-free expression and nanodisc integration of RferOR1

The open reading frame (ORF) of RferOR1 [13] was cloned into the pET20b+ vector under the control of a T7 promoter, enabling its expression in a cell-free system in accordance with previously described protocols for RferOR1 cell-free expression [27]. We expressed RferOR1 in an *E. coli*-based cell-free expression system with a 6 His-tag at the N-terminus, by Cube Biotech GmbH (Monheim, Germany). To facilitate detection and purification, a Rho1D4 tag sequence (TETSQVAPA) and a flexible linker (GSSG) were appended to the C-terminus of the RferOR1 protein. The Rho1D4 affinity tag system was subsequently utilized for efficient purification of the expressed RferOR1 proteins.

Expression and solubilization of RferOR1 were carried out in the presence of various detergents, including Brij 35, Brij 58, and n-decyl-β-D-maltoside (DM), at concentrations of 0.4%, 0.2%, and 0.1%. Of the detergents tested, Brij 58 proved to be the most effective for solubilizing the protein. Following solubilization, RferOR1 was purified and prepared for integration into nanodiscs (MSP1D1 and MSP1E3D1) (Cube Biotech GmbH, Germany). Analysis of the purified protein fractions was performed using Western blotting with both anti-His and anti-Rho1D4 antibodies, as well as by Coomassie-stained gel electrophoresis, to confirm the presence and purity of the RferOR1 protein.

The RferOR1 nanodisc integration was outsourced to Cube Biotech (Monheim, Germany). Two distinct nanodisc scaffold proteins were employed for assembly: MSP1D1, which forms nanodiscs of approximately 8 nm in diameter, and MSP1E3D1, which yields nanodiscs of about 13 nm in diameter. The size and homogeneity of the assembled nanodiscs were confirmed using Dynamic Light Scattering (DLS). During the assembly process, bio-beads played a crucial role by removing excess detergents and lipids from the mixture, thereby forming empty nanodiscs suitable for subsequent protein integration.

### 2.9 Biosensor development

OBPs and ORs were immobilized on 20 MHz QCMs using self-assembled monolayers [17, 28]. Pristine sensors were cleaned with dichloromethane before lipid monolayer formation. Saturated vapors were generated by adding ∼20 μL of analyte to a 40 mL vial, sealing it, and leaving it at room temperature for 1–2 hours.

#### 2.9.1 Electronic nose

The electronic nose platform was designed to incorporate up to twelve Quartz Crystal Microbalance (QCM) sensors, all contained within a compact cell measuring 8 cm³. To ensure accurate environmental monitoring, the platform was equipped with integrated temperature and humidity sensors (SHT31, Sensirion, Switzerland). Each QCM sensor is individually connected to its own oscillator circuit. High-precision frequency measurement is achieved by means of a thermally stabilized reference oscillator, enabling precise measurement of its output frequency. These frequencies are determined using a temperature-compensated reference quartz crystal, which provides high resolution down to 0.1 Hz [29].

The electronic system is implemented on a Field Programmable Gate Array (FPGA), delivering robust, flexible signal processing capabilities. Gas sampling and flow are managed by an embedded miniature diaphragm pump—low noise, high precision, and durable—capable of delivering flow rates between 0 and 200 standard cubic centimeters per minute (sccm). For optimal gas distribution, an optimised microchannel manifold, fabricated from polymethyl methacrylate (PMMA) using Computer Numerical Control (CNC) machining, is employed. This arrangement ensures reliable and reproducible delivery of gas samples to the sensor array for subsequent analysis.

For laboratory measurements, CaCl₂ dried ambient air served as the carrier gas for headspace extraction, vial repressurization, sensor baseline recovery, and reference measurements. A two-way valve alternated between reference and sample streams. Each measurement involved a 40-second sample exposure with airflow held at 50 sccm. The system operated *via* USB power, with field tests using a power bank for over 4 hours of continuous use. Data acquisition was managed through custom MATLAB software.

### 2.10 Data analysis

To evaluate differences in sensor signal responses to the various compounds tested, the Kruskal-Wallis rank-sum test was applied. This nonparametric statistical method enabled the assessment of whether significant differences existed among sensor response groups, providing an initial overview of the dataset’s structure. Following statistical evaluation, the sensor signals were organized into matrices for further multivariate analysis. Principal Component Analysis (PCA) and Linear Discriminant Analysis (LDA) were applied to explore and classify the data. Prior to these analyses, standardization was performed on each variable within the matrices, ensuring that every variable had a mean of zero and a variance of one. This step was essential for variable comparability and to optimize the performance of subsequent analyses.

The LDA classifiers were optimized using k-fold cross-validation, a method that partitions the data into k subsets to evaluate the model’s predictive performance rigorously. To assess classification effectiveness, standard performance metrics were calculated, including accuracy, sensitivity, specificity, and the area under the Receiver Operating Characteristic (ROC) curve. These metrics provided a comprehensive evaluation of the model’s ability to identify and distinguish analytes from sensor signals correctly.

## 3. Results

### 3.1 In silico mutagenesis and docking screening

The amino acid sequences of two previously identified R. ferrugineus odorant-binding proteins, RferOBP1768 and RferOBP23 [15], were used as templates for three-dimensional structure modelling using the I-TASSER platform (Figure S1). Binding pocket composition was characterised using CASTp analysis. For RferOBP1768, 33 of 111 residues were identified as constituents of the primary binding pocket (Table S1); of these, 26 (78.8%) were hydrophobic, 5 (15.1%) were positively charged, and 2 (6.1%) were negatively charged (Table S2). For RferOBP23, 17 of 122 residues formed the primary binding pocket (Table S3); of these, 13 (76.4%) were hydrophobic, 2 (11.8%) were positively charged, and 2 (11.8%) were negatively charged.

Thermodynamic stability analysis [delta energy (ΔΔG)] *via* direct amino acid substitution using the PoPMuSiC algorithm analysis [17] identified 21 potentially stabilising mutations (from 627 possible substitutions) at seven distinct binding pocket positions in RferOBP1768 (Table S2); all introduced residues were hydrophobic (Table S3). For RferOBP23, only 2 potentially stabilising mutations were identified from 323 possible substitutions at two binding pocket positions, both introducing hydrophobic residues (Table S2).

Virtual docking was subsequently performed with the *R. ferrugineus* aggregation pheromone components, ferrugineol and ferrugineone, as well as VOCs emitted by healthy and herbivore-damaged palm trees. The docking results were visualised using the Swissdock plugin UCFS Chimera (Figure S2), and the data are presented in (Supplementary Appendix 1). For RferOBP1768, six mutant proteins exhibiting higher predicted binding affinities than wild-type (WT) for the selected ligands arose from substitutions at two residues — Q4 and Q12 — both of which are hydrophilic rather than hydrophobic (Table S4), suggesting that these positions are tolerant of substitution without compromising the global protein fold. For RferOBP23, one of the two stable mutants exhibited higher predicted binding affinities than WT for the selected ligands, arising from substitution of the positively charged residue R49. For both proteins, mutant variants with the highest predicted binding affinities introduced hydrophobic residues at the substitution sites (Tables S3 and S4), consistent with the well-documented hydrophobic character of OBP and OR binding pockets [13, 17].

### 3.2 Protein expression and characterisation

Based on the docking screening results (Supplementary Appendix 1), two OBP mutants — RferOBP1768_Q12V and RferOBP23_R49L — together with the corresponding WT proteins were selected for expression. All proteins were produced with an N-terminal 6×His tag to facilitate purification (Figure S3). Proteins were expressed and purified commercially; purity and identity were confirmed by SDS-PAGE and western blotting (Figures S3 and S4). Ligand-binding affinities were characterised using a fluorescence competitive displacement assay [17] against the target ligands (Supplementary Appendix 1).

### 3.3 Comparison of theoretical and experimental binding data

Experimentally determined binding constants confirmed that WT RferOBP1768 exhibits superior binding affinity (1/K∼D∼) for all ligands tested relative to the other proteins examined, apart from ferrugineone (Table S5). Specifically, WT RferOBP1768 displayed 2.3-fold and 4.8-fold higher affinities compared to WT RferOBP23 for ferrugineol and ethyl acetate, respectively, while WT RferOBP23 exhibited the highest affinity for ferrugineone across all proteins tested. These results are consistent with the computational docking data (Supplementary Appendix 1).

Structural analysis provides a mechanistic basis for the superior binding performance of RferOBP1768. Compared to RferOBP23, RferOBP1768 possesses a binding pocket with approximately 3-fold greater molecular surface area and 3-fold greater pocket depth (Table S3). Furthermore, the binding pocket entrance of RferOBP1768 is wider by 6.15 Å² in mouth surface area and 6.29 Å in mouth length relative to RferOBP23 (Table S3). These structural differences collectively suggest that target ligand molecules can penetrate more deeply into the RferOBP1768 binding pocket (Figure 1), resulting in higher binding affinities compared to RferOBP23 (Figure 1B) — a phenomenon previously reported for OBPs in mosquito species [17].

**Figure 1.**
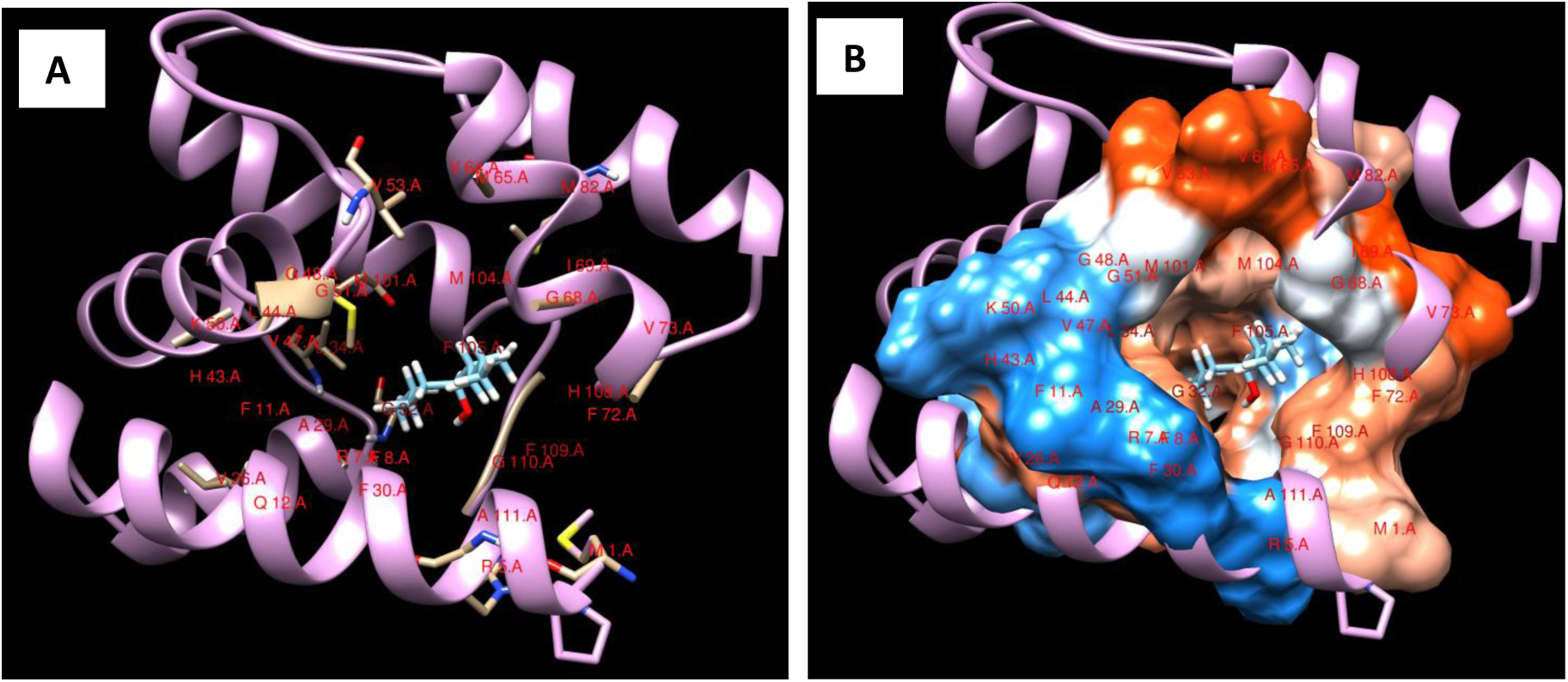
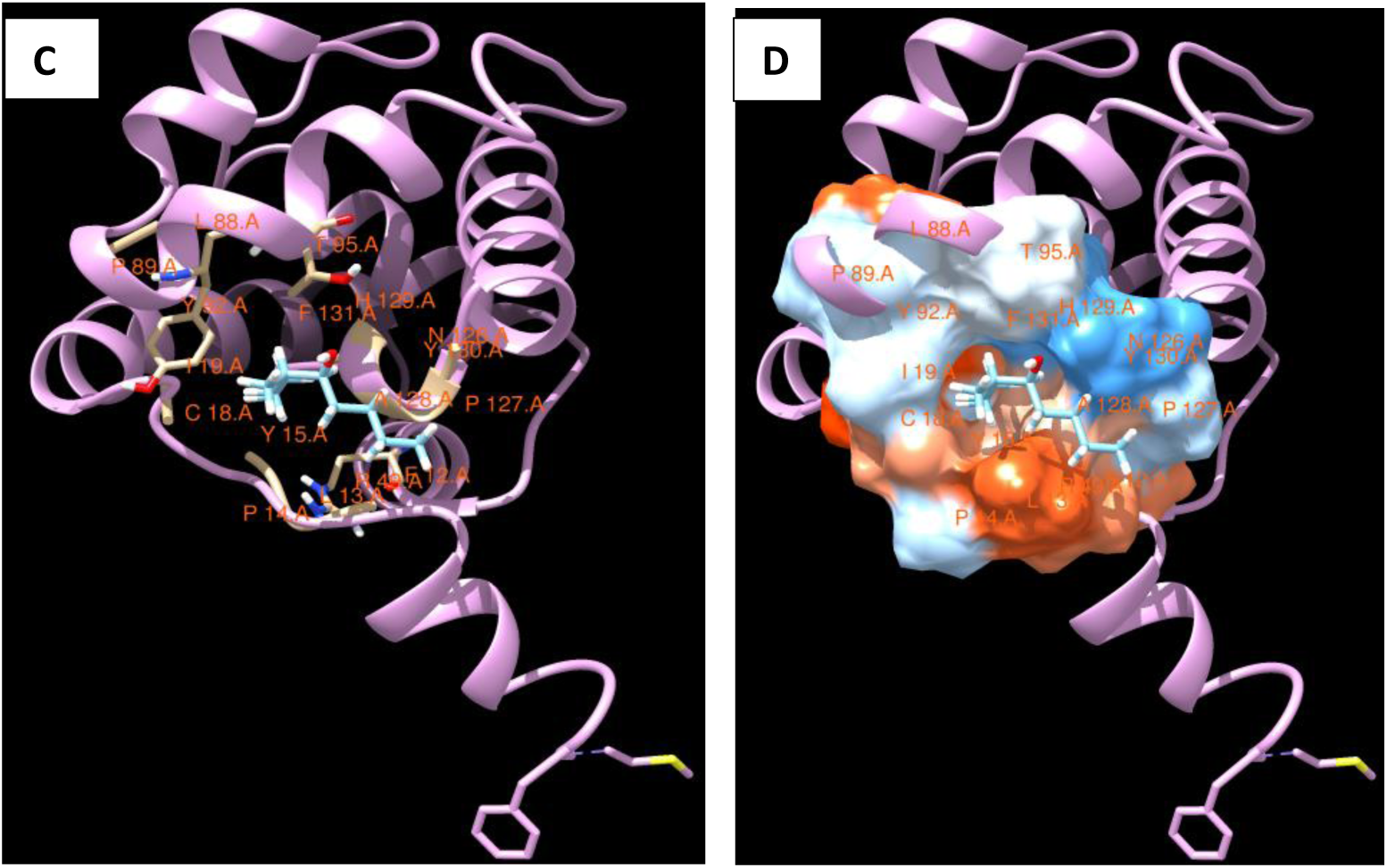
Binding of ferrugineol to the primary binding site of: (A, B) WT RferOBP1768 and (C, D) WT RferOBP23. (A) Cartoon representation and (B) hydrophobicity surface representation of the binding pocket of RferOBP1768; ferrugineol docks fully within the binding pocket with a binding energy of −33.99 kcal/mol. (C) Cartoon representation and (D) hydrophobicity surface representation of the binding pocket of RferOBP23; ferrugineol does not fully accommodate within the shallower binding pocket, yielding a binding energy of −27.97 kcal/mol. Lower binding energy corresponds to higher predicted binding affinity. Binding pocket residues are highlighted in red.

Experimentally determined binding affinities of mutant OBPs were generally reduced relative to the WT proteins, with a few exceptions, contrary to predictions from the docking screening (Table S5). Two factors likely account for this discrepancy. First, ligands may access the mutant binding pockets via alternative binding routes rather than the canonical pathway operative in WT proteins [17]; the docking algorithm preferentially ranks binding modes within the canonical pocket, assigning lower energy values — corresponding to higher predicted affinities — to these configurations, which may not accurately reflect the altered binding landscape of mutant proteins. Second, the mutant OBPs displayed reduced binding affinities for the fluorescence displacement probe 1-NPN relative to WT proteins (Table S5); since the apparent affinity of competing ligands in this assay is proportional to probe affinity, reduced probe binding in the mutant proteins will systematically underestimate the apparent affinity of competing ligands. Importantly, the rank order of experimentally measured binding constants was consistent with the computational predictions, validating the overall in silico prioritisation strategy.

Despite the reduced absolute affinities of several mutant proteins, the combination of WT and mutant OBPs exhibiting a diverse range of binding affinities across the tested ligand panel is strategically advantageous for sensor array design. The differential binding profiles of individual array elements generate distinct, partially overlapping response patterns that — in direct analogy with the combinatorial coding principle of biological olfaction [30] — provide the discriminatory power required to distinguish between structurally similar target analytes in complex volatile backgrounds.

Approximately 0.55 mg of RferOR1 was produced in Brij-58 detergent (Figure S4), concentrated, and subjected to nanodisc integration. RferOR1 was reconstituted into lipid nanodiscs of two distinct sizes: MSP1D1 (8 nm diameter, 26 kDa; 0.44 mg/ml, ∼230 μl) and MSP1E3D1 (13 nm diameter, 31 kDa; 0.64 mg/ml, ∼240 μl) (Figure S5), referred to hereafter as OR1_1_ and OR1_2_, respectively, to assess the effect of nanodisc size on receptor performance. Further, an ultrafiltration centrifugal concentrator concentrated the OR1. The quality and characteristics of the resulting nanodiscs were assessed using dynamic light scattering (DLS) measurements, which confirmed a single sharp peak for uniform samples (Figure S5). The polydispersity index (PDI) obtained (0.1 to 0.2) was found acceptable for standard, good-quality nanodisc preparations.

### 3.4 Biosensor fabrication

To achieve vapour-phase analyte detection, proteins (RferOBP1768, RferOBP23 and RferOR1) were immobilised onto 20 MHz quartz crystal microbalance (QCM) transducers via self-assembled monolayer (SAM) chemistry [17]. The functionalised QCM biosensors were assembled into a sealed sensor cell and connected to dedicated electronic circuitry for resonant frequency measurement and data acquisition. In QCM-based sensing, a change in mass (Δm) at the quartz surface produces a proportional shift in oscillation frequency (Δf), enabling label-free, real-time detection of molecular binding events [29]. All functionalised sensors responded sensitively to vapour-phase exposures of the tested analytes (Figure 2).

**Figure 2.**
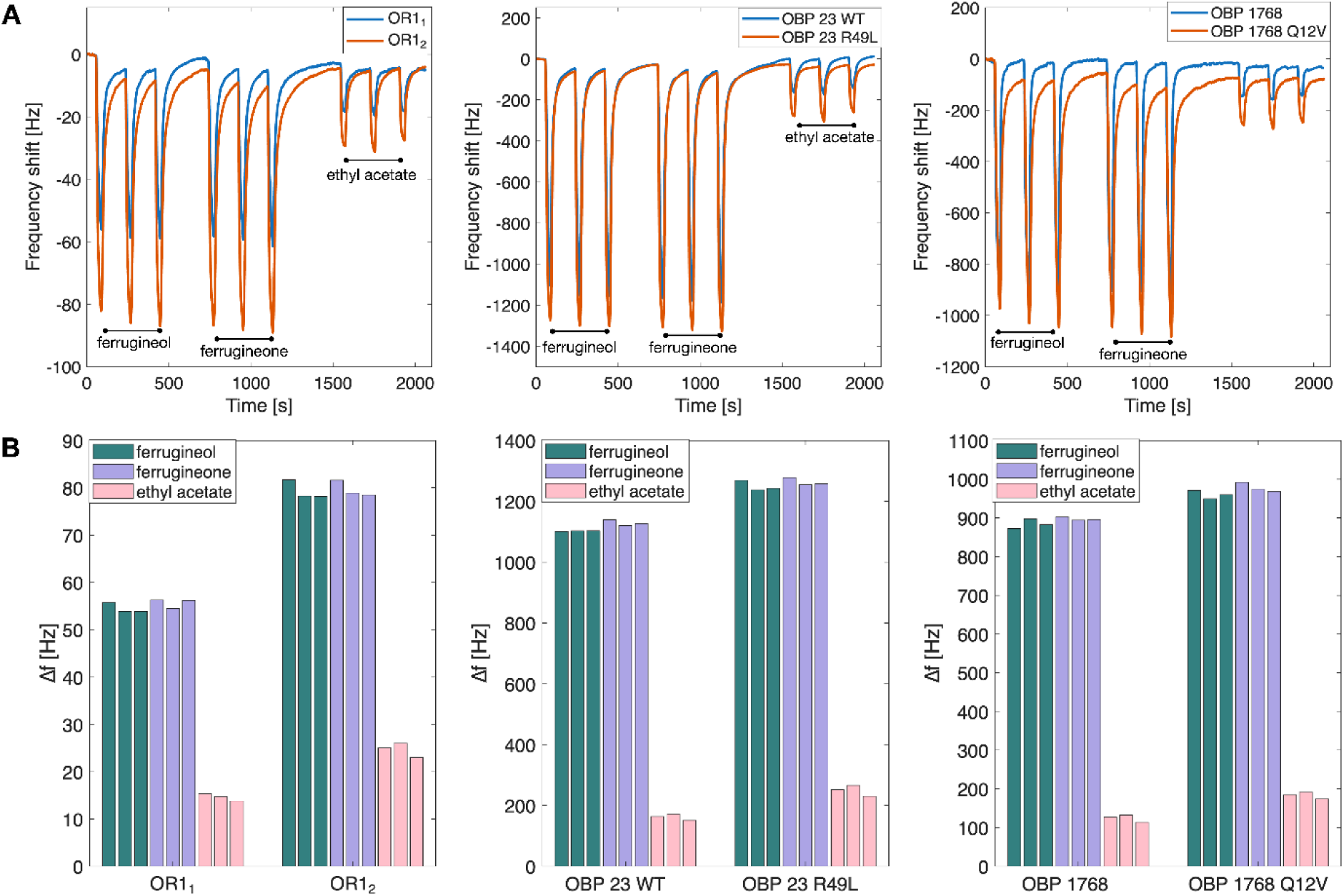
(A) Raw sensor response signals recorded during three consecutive saturated vapour pulses of ferrugineol, ferrugineone, and ethyl acetate. Responses of WT and mutant OBPs and RferOR1 in nanodiscs of two sizes are compared. (B) Net frequency shift, calculated as the difference between baseline frequency immediately before and at the end of each vapour pulse, representing the sensor response magnitude. OR1 sensor responses vary with nanodisc size. Mutant OBPs consistently produce slightly greater responses than the corresponding WT proteins.

### 3.5 Sensor characterization

The experimental setup for sensor characterisation is described in Figure S6. Sensors were supplied with a continuous stream of synthetic air to establish a stable baseline prior to all measurements. Sensor responses were evaluated against a panel of volatile compounds known to elicit electrophysiological responses in *R. ferrugineus* antennae, including *R. ferrugineus* aggregation pheromones, structurally homologous alcohols and ketones with documented OBP interactions [12–14], aggregation pheromones from related non-sympatric weevil species, and palm-emitted host plant volatiles acting as kairomonal attractants (Table 1).

For each measurement, the airflow was transiently diverted through a bubbler containing the test compound in liquid form, generating saturated vapour for approximately 30 seconds, before returning to synthetic air. Figure 2A shows sensor signals recorded during repeated exposures to saturated vapours of ferrugineol, ferrugineone, and ethyl acetate. Sensor signals returned consistently to baseline after each exposure, confirming the reversibility of analyte–protein interactions — a prerequisite for reusable sensor operation.

The raw sensor response was calculated as the net frequency shift between the baseline signal immediately before and at the end of each vapour pulse (Figure 2B). Response magnitude depends on both the surface density of immobilised protein on the QCM electrode and the intrinsic binding affinity of the protein–ligand interaction. The comparatively lower responses of RferOR1 sensors relative to OBP sensors likely reflect lower receptor surface coverage on the gold electrode, attributable in part to the larger physical dimensions of the nanodisc–receptor complexes. The dependence of RferOR1 sensor response magnitude on nanodisc size is consistent with this interpretation, while mutant OBPs consistently outperformed their WT counterparts in response magnitude.

Direct comparison of sensor responses requires normalisation to account for the substantial differences in compound vapour pressure among the tested analytes (Table 1). For example, ethyl acetate has a saturated vapour pressure approximately 1,330-fold higher than ferrugineol and approximately 200-fold higher than ferrugineone. Despite this large concentration variability, sensor responses to the *R. ferrugineus* pheromones were 4–6-fold greater than responses to ethyl acetate (Figure 2), demonstrating the selectivity of RferOR1 and OBPs for their cognate pheromone ligands. All sensors produced measurable responses to all tested compounds. This broad but selective response profile is consistent with single-sensillum recordings (SSR) from transgenic *Drosophila* expressing RferOR1, which similarly showed preferential but not exclusive responses to *R. ferrugineus* pheromone components [13, 14].

Sensor responses to pheromones and structurally related interfering compounds were measured following exposure to the corresponding saturated vapour pressures, with all measurements performed in triplicate. The measurement sequence was randomised to minimise hysteresis effects. Given the large differences in saturated vapour pressures among test compounds (Table 1), raw sensor responses were normalised by dividing the frequency shift by the corresponding saturated vapour pressure, yielding a quantity proportional to the apparent sensor affinity for each compound, independent of analyte concentration.

Figure 3 presents the vapour-pressure-normalised sensor responses as box plots. All sensors displayed marked affinity for ferrugineol, while affinity for ferrugineone — the minor *R. ferrugineus* pheromone component — was generally lower. Notably, the mutant OBPs RferOBP23_R49L and RferOBP1768_Q12V exhibited significantly higher normalised affinities for ferrugineone relative to their respective WT proteins, demonstrating that targeted mutagenesis can shift ligand selectivity within the pheromone component panel. RferOR1 sensors produced qualitatively similar response patterns regardless of nanodisc size (OR1_1_ versus OR1_2_), differing principally in absolute response magnitude (Figure 2B).

**Figure 3.**
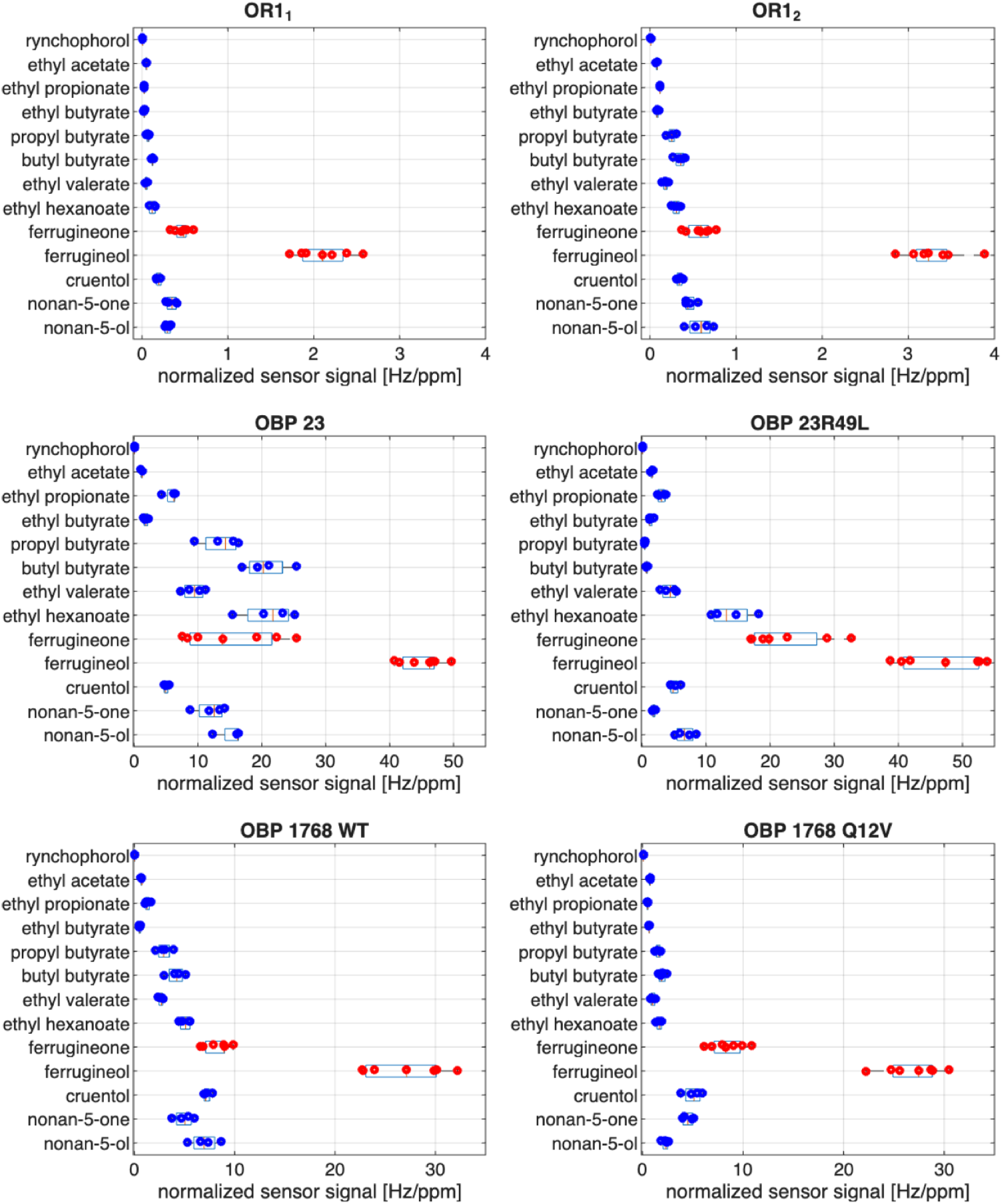
Normalised sensor response profiles of the six OBP- and OR-functionalised sensors to the full panel of test volatile compounds. Each compound was measured at minimum in triplicate. Sensor responses were normalised by the saturated vapour pressure of each compound (Table 1) and are presented as box plots illustrating the statistical variability of replicated measurements. *R. ferrugineus* pheromone components are highlighted in red.

### 3.6 Sensor array performance and analyte discrimination

The normalised response profiles in Figure 3 demonstrate that no individual sensor can reliably discriminate between *R. ferrugineus* pheromone components (ferrugineol and ferrugineone) and structurally related plant-emitted esters under realistic field conditions, where palm-derived VOCs are present at substantially higher concentrations than pheromones [12]. Discrimination of infested from non-infested palms therefore requires a sensor array exploiting combinatorial selectivity — the principle that cross-reactive sensors with distinct but partially overlapping specificities collectively generate high-dimensional response patterns sufficient for analyte classification [31].

To identify the minimal sensor array configuration sufficient for complete analyte discrimination, normalised responses from all six sensors were evaluated in all possible array configurations (63 permutations). For each configuration, a Linear Discriminant Analysis (LDA) classifier was trained and tested on standardised data to eliminate intensity bias, and classification accuracy (percentage of correct identifications) was determined. As shown in Figure 4A, classification accuracy increased monotonically with array size, though the rate of improvement depended on array composition. The optimal sensor combination for each array dimension was identified by exhaustive permutation and is represented as a heat map in Figure 4B.

**Figure 4.**
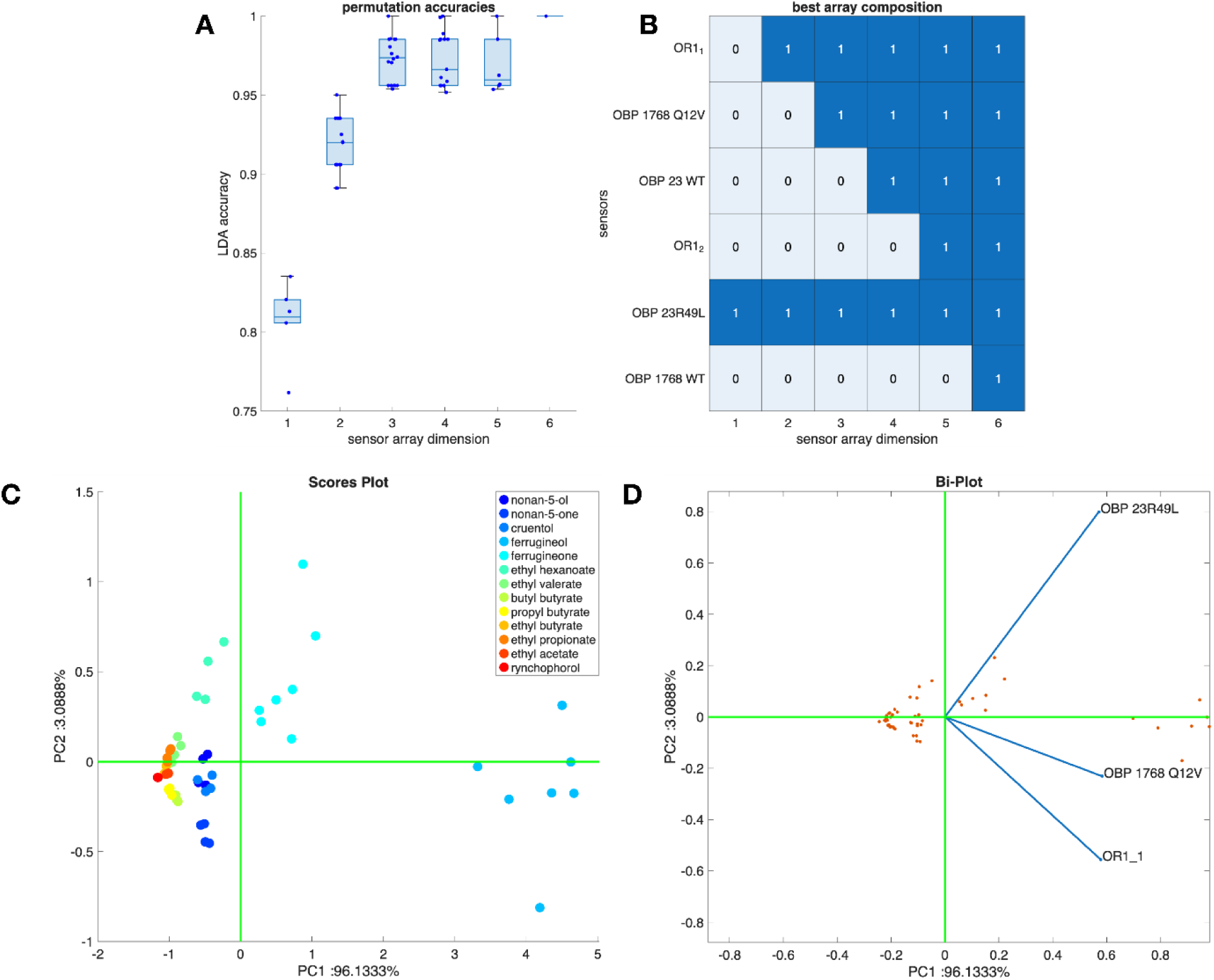
Sensor array classification performance. (A) Classification accuracy of LDA models as a function of array dimension; for each dimension, the distribution of accuracy across all possible array compositions is shown as a box plot. (B) Optimal array composition for each dimension represented as a heat map. (C) PCA scores plot of the optimal three-sensor array. (D) Corresponding biplot showing sensor loadings overlaid on the scores plot.

The minimal array achieving 100% classification accuracy comprised two mutant OBPs (RferOBP23_R49L and RferOBP1768_Q12V) and RferOR1 reconstituted in the smaller MSP1D1 nanodisc (OR11). Incremental addition of sensors demonstrated a stepwise improvement in accuracy: RferOBP23_R49L alone achieved 83% accuracy; addition of OR11 increased accuracy to 95%; and incorporation of RferOBP1768_Q12V achieved 100%. Notably, RferOR1 in the larger MSP1E3D1 nanodisc (OR1_2_) appeared only in the optimal five-sensor array, suggesting that the increased non-specific van der Waals interactions associated with the larger nanodisc reduce selectivity and diminish its contribution to discrimination at lower array dimensions, despite the higher signal intensity produced (Figure 2B).

Principal Component Analysis (PCA) was applied to the optimal three-sensor array to visualise the discriminatory properties of the minimal configuration (Figure 4C). The first principal component explained 96% of total variance, indicating strong consensus among the three sensors in their response to ferrugineol and confirming that ferrugineol forms a well-resolved cluster, well-separated from all other analytes. Ferrugineone was positioned at the periphery of the homologous compound cluster, reflecting its structural similarity to related ketones. The loading plot (Figure 4D) confirmed that all three sensors contribute to ferrugineol recognition, while RferOR1 and RferOBP23_R49L provide the primary discriminatory power for ferrugineone identification. These results demonstrate that the strategic integration of proteins with complementary and partially overlapping binding profiles generates a multidimensional response space that enables robust discrimination of target pheromone components from complex volatile backgrounds.

### 3.7 Sensor validation on real samples

The optimal sensor array — comprising mutant OBPs and RferOR1 — was validated against authentic volatile profiles from live insects and palm trees (Figure 5). Sensors were integrated into the electronic nose platform used for laboratory characterisation, equipped with an air-sampling system. For all real-sample measurements, sensor responses were calculated as the difference between the sample signal and a clean ambient air baseline obtained through activated carbon filtration.

**Figure 5.**
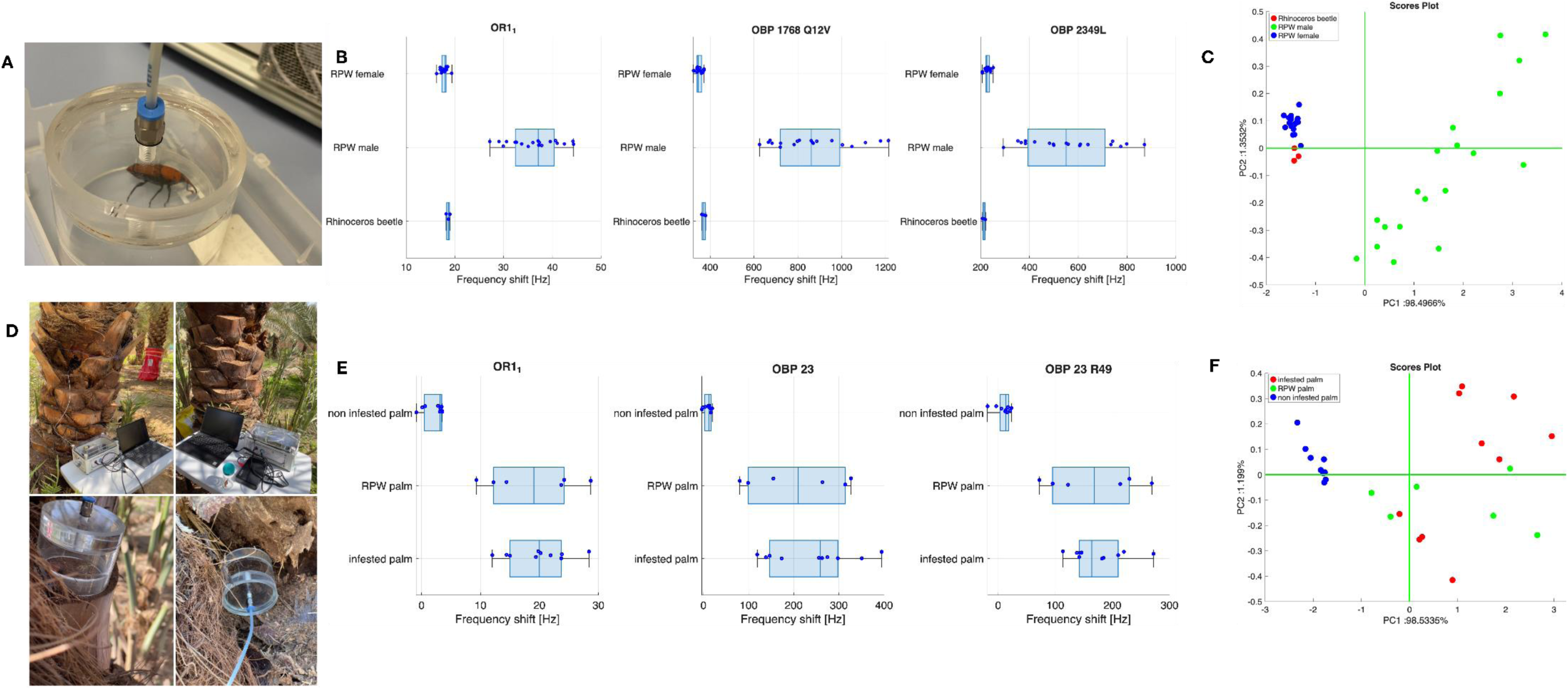
Real-sample and field validation. (A) Schematic of the insect headspace volatile sampling chamber. (B) Sensor array responses to volatile emissions from male *R. ferrugineus*, female *R. ferrugineus*, and *O. rhinoceros*. (C) PCA scores plot of data shown in (B). (D) In-field volatile sampling setup for palm trees and field-collected R. ferrugineus adults; a sealed plastic cup defines the sampling volume, and the electronic nose — controlled and powered by a laptop computer — draws headspace volatiles across the sensor array. (E) Sensor array responses to infested palms, non-infested palms, and field-collected *R. ferrugineus* adults. (F) PCA scores plot of data shown in (E).

### 3.8 Laboratory validation using live insects

Volatile emissions from laboratory-cultured male and female *R. ferrugineus* and from the coconut rhinoceros beetle, *Oryctes rhinoceros* — a species that co-inhabits palm field environments with *R. ferrugineus* but produces a chemically distinct aggregation pheromone, ethyl 4-methyloctanoate [32].

Sensor responses (Figure 5B) were consistently and substantially higher for male *R. ferrugineus* than for female *R. ferrugineus* or *O. rhinoceros*. Given that all insects were maintained on identical diets and under equivalent rearing conditions, this differential response is most parsimoniously attributed to the male-specific production and release of *R. ferrugineus* aggregation pheromones [10]. Responses to female *R. ferrugineus* and *O. rhinoceros* were of comparable magnitude, suggesting that non-pheromonal background volatiles elicit weaker and less specific sensor binding. Inter-individual variability in male *R. ferrugineus* responses likely reflects natural fluctuations in pheromone biosynthesis and emission rate. PCA of the insect volatile data (Figure 5C) confirmed unambiguous discrimination of male *R. ferrugineus* from the other insect species and revealed a consistent, albeit more subtle, separation between female *R. ferrugineus* and *O. rhinoceros*, indicating that the sensor array retains residual discriminatory sensitivity to non-pheromonal species-specific volatiles beyond the primary pheromone signal — a detection feature amplified by the mass-sensitive QCM platform, which responds to all molecular binding events at the transducer surface.

### 3.9 Field validation on palm trees

Field trials were conducted in an experimental date palm orchard in Riyadh, Saudi Arabia, where individual trees were under continuous surveillance for *R. ferrugineus* infestation. Headspace volatile sampling was performed non-invasively by positioning a sealed plastic sampling cavity against the palm trunk bark surface (Figure 5D). Sensors were also exposed to individual *R. ferrugineus* adults encountered on tree surfaces during the field campaign and temporarily retained for direct measurement.

Sensor responses to field-collected *R. ferrugineus* adults and to confirmed infested palm trees were similar in magnitude and both significantly higher than responses recorded for non-infested trees (Figure 5E). PCA of the field data (Figure 5F) revealed a clear and statistically robust separation between infested and non-infested palms, with responses from on-tree *R. ferrugineus* adults clustering with those of infested palm trees. This clustering strongly suggests that the volatile signals detected in infested palm headspace originate predominantly from aggregation pheromones released by resident weevils or accumulated within infested palm tissue, rather than from plant-derived stress volatiles. Inter-tree variability among infested palms likely reflects natural differences in infestation severity, weevil population density, and pheromone emission dynamics at the time of sampling. Collectively, these field results confirm that the sensor array reliably detects *R. ferrugineus* male aggregation pheromones under authentic field conditions and, through residual affinity to species-specific non-pheromonal volatiles, can discriminate infested from non-infested palms and distinguish *R. ferrugineus* from co-occurring non-target insect species.

### 3.10 Biosensor operational stability

The long-term operational stability of immobilised OBP- and OR-functionalised QCM biosensors was assessed by monitoring sensor responses to periodic pulses of saturated ferrugineol vapour over extended storage periods under ambient conditions. OBP-functionalised sensors (RferOBP1768 and RferOBP23) retained full functional activity across the entire 12-month monitoring period (Figure 6A), demonstrating exceptional long-term stability unprecedented for protein-based biosensors. The RferOR1 QCM biosensor remained fully functional and produced reproducible responses to ferrugineol vapour over a continuous 220-day (approximately 7-month) monitoring period (Figure 6B), substantially exceeding the operational stability of any previously reported odorant receptor-based biosensor. Day-to-day variability in response magnitude was attributed to fluctuations in saturated vapour concentration generated under ambient temperature conditions [17], rather than to sensor degradation.

**Figure 6.**
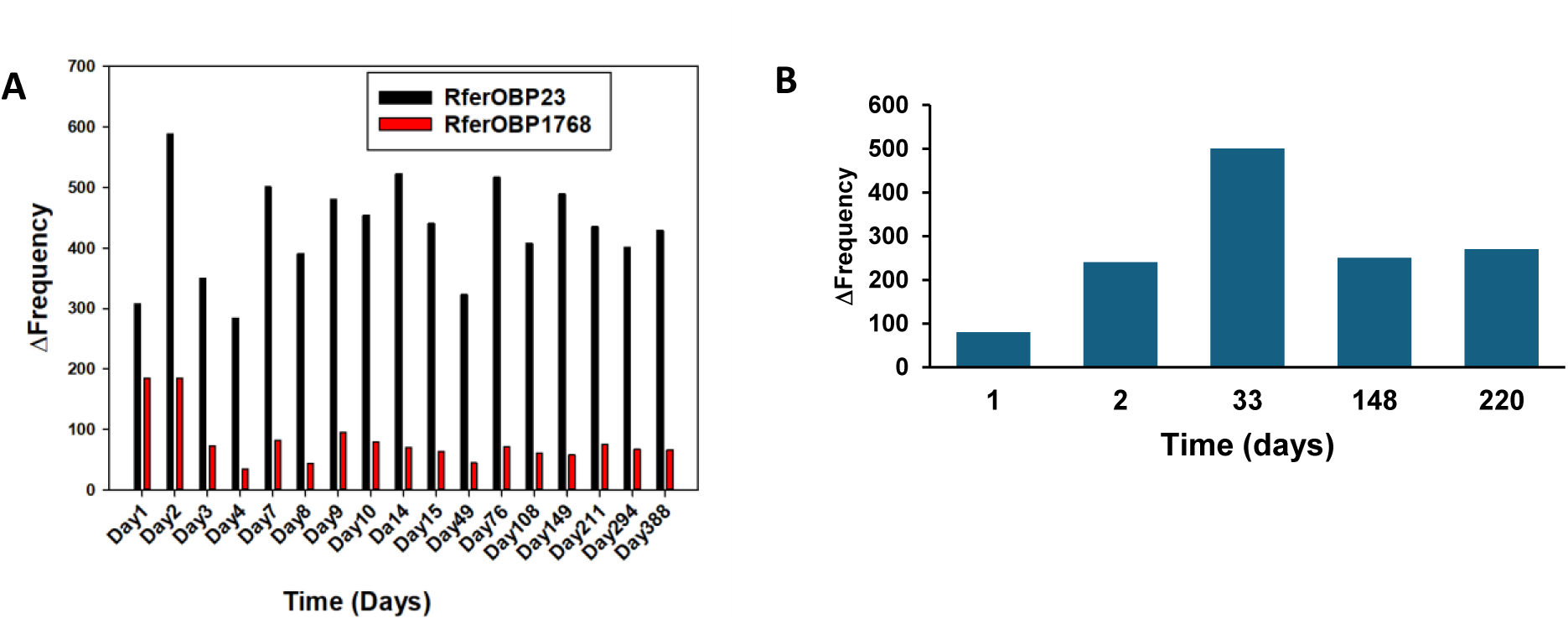
Long-term operational stability of immobilised biosensors assessed by repeated exposure to saturated ferrugineol vapour. (A) RferOBP23 and RferOBP1768 biosensor stability monitored over 12 months. (B) RferOR1 biosensor stability monitored over 220 days (∼7 months). Response variability reflects day-to-day fluctuations in ambient-temperature saturated vapour concentration rather than sensor degradation.

## 4. Discussion

In this study, we developed a biohybrid sensor array capable of discriminating between weevil-infested and healthy date palms at early stages of infestation. The array integrates a pheromone receptor (RferOR1) stabilised within lipid nanodiscs and odorant-binding proteins (RferOBP1768 and RferOBP23), both immobilised on quartz crystal microbalance (QCM) transducers. Whereas previous OR-based biosensors have exclusively detected odorants in the liquid phase [21, 22, 33], we demonstrate for the first time that OR- and OBP-functionalised QCM transducers can detect airborne odorants, enabling the development of robust, field-deployable detection devices. The integration of ORs and OBPs enabled the biosensor array to functionally mimic the weevil antenna, achieving precise detection of the *R. ferrugineus* aggregation pheromone within complex volatile blends present in palm plantation environments. The sensor array responded sensitively to low analyte concentrations and discriminated RPW-specific pheromone components from volatiles produced by other weevil species and from volatiles emitted by abiotic stress-challenged palm trees. Importantly, both OR- and OBP-functionalised sensors retained full functional activity for at least seven months under ambient storage conditions, demonstrating the feasibility of long-term, low-maintenance field monitoring. While numerous biosensor technologies have been reported in recent years [8, 21, 34], including a cell-based biosensor in *R. ferrugineus* [9], to our knowledge the present biohybrid array is the first to achieve early-stage palm weevil detection under genuine field conditions, representing a significant advance in sustainable agricultural pest surveillance. Notably, the cell-based biosensor reported recently for *R. ferrugineus* early detection has a major limitation: a lack of field trial data, and it has been demonstrated only in lab trials [9].

Palm trees underpin global sustainability, serving as critical food sources and holding immense sociocultural, touristic, and historical value across multiple continents [35]. They sustain hundreds of millions of people by providing food, beverages, fibre, medicine, and shelter, while also supporting biodiversity and sequestering atmospheric carbon, thereby contributing to climate change mitigation [36]. These ecological and economic services are increasingly threatened by the invasion of *R. ferrugineus* across Asia and the Mediterranean and *R. palmarum* across the Americas, both of which cause severe, often irreversible damage to palm plantations and undermine their ecological and economic value [2, 37]. With growing global concern over palm tree sustainability, early detection of weevil infestation is an urgent priority [2], as timely intervention enables targeted, ecologically informed management to protect crop yield, quality, and food security. Unlike previously reported cell-based biosensors demonstrated exclusively under laboratory conditions [9], the present biohybrid sensor array is specifically engineered for the detection of airborne aggregation pheromones released by *R. ferrugineus* in infested palm plantation environments, and has been validated under genuine field conditions for early detection and real-time monitoring of weevil infestation.

Functionalising sensors with OBPs and ORs is a well-recognised strategy for constructing artificial olfactory systems [38]. OBPs are structurally compact, thermostable, and highly resilient to environmental stressors including temperature fluctuations and organic solvents [39]. OBP-functionalised sensors retain the intrinsic ligand sensitivity and selectivity of the native protein [40, 41], and OBP-QCM devices have demonstrated practical utility in demanding real-world applications, including search-and-rescue operations [42]. Integration of ORs into biosensor platforms is considerably more challenging owing to their dependence on a lipid bilayer membrane environment and their requirement for the obligate co-receptor Orco, which forms a ligand-gated heteromeric ion channel with OR subunits [20]. Recent studies have reconstituted functional OR–Orco complexes in lipid bilayer systems for the detection of volatile organic compounds [43]. In contrast, our approach circumvented the requirement for Orco co-expression by stabilising RferOR1 within lipid nanodiscs, substantially simplifying sensor fabrication. QCMs provided a robust and versatile transducer platform; however, their sensitivity to weak non-specific interactions with airborne molecules presents a potential challenge to selectivity. To mitigate non-specific binding and enhance discrimination capacity, we applied the combinatorial coding principle of biological olfaction [44] by integrating multiple QCM sensors with differing binding profiles into an array, enabling pattern-based analyte identification analogous to the processing of electronic nose data.

A central challenge in OR-based sensor design is maintaining the structural integrity and ligand-binding activity of membrane-embedded receptor proteins outside their native lipid environment. Nanodisc technology provides a defined, stable lipid scaffold that preserves membrane protein conformation and has previously been employed in biosensor applications using human and *Drosophila* ORs [45]. In the present study, RferOR1 embedded in lipid nanodiscs retained full pheromone detection capability, and both OR- and OBP-based sensor components retained functionality outside aqueous environments. Nanodisc stabilisation ensured conformational integrity of RferOR1, enabling parts-per-million-level sensitivity for airborne pheromone detection under field conditions in palm plantations.

*In silico* mutagenesis was employed to characterise and engineer the ligand-binding pockets of OBPs, enabling rational selection of protein variants with tailored binding affinities for target volatile compounds [17]. Virtual docking of target ligands to computationally generated mutant OBPs facilitated pre-selection of candidate variants for experimental expression and binding characterisation, substantially reducing empirical screening effort. A key methodological challenge in this approach was establishing an appropriate energy threshold to define whether a given mutation improved or diminished ligand binding relative to the wild-type protein; accordingly, the threshold applied was empirically calibrated. Experimentally measured binding affinity constants deviated quantitatively from values predicted by molecular docking — a recognised limitation arising from the inherent assumptions of computational modelling tools and from the fluorescence displacement assay employed to determine experimental dissociation constants. Nevertheless, the rank order of experimentally measured binding constants was consistent with the computational predictions, validating the *in silico* prioritisation strategy. Wild-type and mutant OBPs exhibited a range of experimentally determined binding affinities for the target ligands, with each variant displaying a distinctive binding activity profile — collectively providing a broad spectrum of differential selectivity. This diversity of binding profiles is directly exploitable in a sensor array context: the combinatorial pattern of binding affinities across array elements provides sufficient discriminatory power to distinguish between structurally similar analytes, in direct analogy with the pattern recognition approach underpinning electronic nose data processing.

Biosensor responses were consistently stronger to RPW-specific pheromone components than to other volatiles tested, and both OR- and OBP-based sensors responded more robustly to volatiles from male *R. ferrugineus* than from females, consistent with the established male-specific release of aggregation pheromones [10]. Field trials confirmed high selectivity for weevil-infested palms over healthy control trees. Limits of detection were estimated from sensor response data and vapour pressure calculations: for ferrugineol (vapour pressure approximately 73 ppm), the measured sensor response of approximately 1,200 Hz yielded, supposing a linear adsorption isotherm, an estimated sensitivity of 16 Hz/ppm, and a nominal noise floor of 1 Hz corresponds to a calculated detection limit of approximately 60 ppb. Applying the same calculation to ferrugineone yields an estimated detection limit of approximately 370 ppb. These detection limits are well within the range of pheromone concentrations encountered in field conditions, supporting practical applicability. It is important to note that, at low concentrations, the adsorption isotherm is expected to exhibit Langmuirian behaviour. Consequently, the linear approximation adopted here may result in an underestimation of the sensor figures of merit.

With respect to operational stability, OBP-functionalised sensors retained binding activity for over 12 months, considerably exceeding the shelf life of comparable biosensor systems reported in the literature. OR-based sensors retained pheromone detection sensitivity for at least seven months, surpassing the longevity of any previously reported OR biosensor. Collectively, these stability data indicate that the OR–OBP array is suitable for deployment in long-term commercial pest surveillance programmes, though further large-scale field validation across diverse palm cultivation environments and climatic conditions is warranted before full commercialisation.

Looking ahead, the odorant-sensing platform developed here is inherently modular and can be adapted for surveillance of other insect pest species once their OBPs and ORs have been functionally characterised [46]. The functional characterisation of RpalOR1 — the *R. ferrugineus* RferOR1 orthologue in *R. palmarum* — has established that both receptors exhibit highly similar response profiles to RPW aggregation pheromone components and structurally related molecules [14], and that *R. ferrugineus* pheromone compounds potently activate RpalOR1. Furthermore, ethyl acetate — a short-chain ester abundantly released by weevil-infested palm tissue — is commonly employed as a co-attractant in pheromone-based mass trapping programmes for both *R. ferrugineus* and *R. palmarum*, and the present biosensor platform can detect ethyl acetate at relevant concentrations. The sensor array therefore offers a practical early detection solution applicable to both palm weevil species. The sensor array therefore offers a practical early detection solution applicable to both palm weevil species. More broadly, the OR–OBP biosensor framework offers a scalable, adaptable foundation for long-term insect pest surveillance that can be expanded to agriculturally important pests beyond palm weevils. It also lends itself to applications in food safety surveillance [47] and hazardous volatile compound detection, capitalizing on the remarkable sensitivity and selectivity of the insect olfactory system as an engineering asset.

## 5. Conclusion

In summary, we successfully fabricated an OR-OBP-sensor that could selectively and sensitively detect the red palm weevil, *Rhynchophorus ferrugineus* pheromone (4RS,5RS)-4-methylnonan-5-ol (ferrugineol). Critically, the sensor array operates without the obligate co-receptor Orco, substantially simplifying device fabrication, as QCM transducers convert molecular binding events directly into measurable resonant frequency shifts without requiring ion channel activity. The array achieved selective detection of airborne ferrugineol at a limit of detection of approximately 60 parts per billion (ppb) under field conditions — well within the range of environmentally relevant pheromone concentrations surrounding infested palms. Field validation in date palm plantations confirmed unambiguous discrimination of weevil-infested from healthy palms, with infested tree volatile profiles clustering with those of live *R. ferrugineus* adults, demonstrating that pheromone signals dominate the infested palm headspace. OBP-functionalised sensors retained full functional activity for over 12 months, and the RferOR1-based sensor remained fully operational for over 7 months under ambient storage conditions, substantially exceeding the operational stability of any previously reported odorant receptor biosensor. Collectively, these results establish a practical, field-validated chemical sensing platform for early-stage *R. ferrugineus* infestation surveillance, offering a sensitive, selective, and sustainable alternative to conventional detection methods for deployment in integrated pest management programmes and precision agriculture. The modular architecture of the OR–OBP sensor array is inherently adaptable to other volatile analyte targets; the recent functional deorphanisation of the *R. palmarum* pheromone receptor RpalOR32 [48] provides a new biorecognition interface for extension of the platform to early monitoring of the South American palm weevil — enabling simultaneous, species-resolved surveillance of both globally destructive palm weevil species from a single integrated sensor array.

## CRediT authorship contribution statement

**Khasim Cali**: Investigation, Formal analysis, Writing – original draft. **Binu Antony**: Conceptualization, Supervision, Investigation, Formal analysis, Writing – original draft. **Corrado Di Natale**: Conceptualization, Supervision, Investigation, Formal analysis, Writing – original draft. **Alexandro Catini**: Investigation. **Nicolas Montagné**: Investigation, Formal analysis. **Emmanuelle Jacquin-Joly**: Conceptualization, Supervision, Writing – review & editing. **Mohammed A. AlSaleh**: Project administration, Field experiment. **Yousef Al-Fehaid**: Field experiment. **Krishna C. Persaud**: Conceptualization, Supervision, Investigation, Formal analysis, Writing – original draft. **Arnab Pain:** Conceptualization, Supervision, Investigation, Writing – review & editing.

## Declaration of generative AI-assisted technologies in the manuscript preparation process

During the preparation of this manuscript, the authors used Claude Sonnet (Claude for free) to improve the English language and readability of the manuscript. After using this tool, the authors reviewed and edited the content as needed and took full responsibility for the content of the published article.

## Acknowledgments

This work was supported by research grants from King Abdullah University of Science and Technology (KAUST), Saudi Arabia (KAUST-OSR-2018-RPW-3816-1, OSR-2018-RPW-3816-4, KAUST-BAS/1/1020-01-01), and the French program “Cultiver et Protéger Autrement” (ANR-20-PCPA-0007). The authors gratefully acknowledge the Deanship of Scientific Research Ongoing Research Funding program – Research Chairs (ORF-RC-2025-3809), King Saud University, Riyadh. We thank the Ministry of Environment, Water and Agriculture (MEWA), Saudi Arabia, for granting access to date palm fields in Al Kharj and Riyadh for field trials.

## Data and materials availability statement

The sensor platforms (devices) developed at the University of Manchester (Persaud lab) and the University of Rome Tor Vergata (Di Natalie’s lab), along with related data acquisition and analysis software, are owned by the respective PIs. RferOR1, RferOBP1768, RferOBP1768_Q12V, RferOBP23_R49L, and RferOBP23 are originally cloned from *R. ferrugineus* antennae and are available at Antony’s lab (King Saud University).

## Supplementary Material

Supplementary material is available Online.

