## Supplementary File: Figures S1 to S6; Tables S1 to S6 for "Insect olfaction-inspired biohybrid sensor array for selective airborne pheromone detection and early monitoring of invasive *Rhynchophorus ferrugineus*"

**Supplementary file includes:**

Figures S1 to S6

Tables S1 to S6

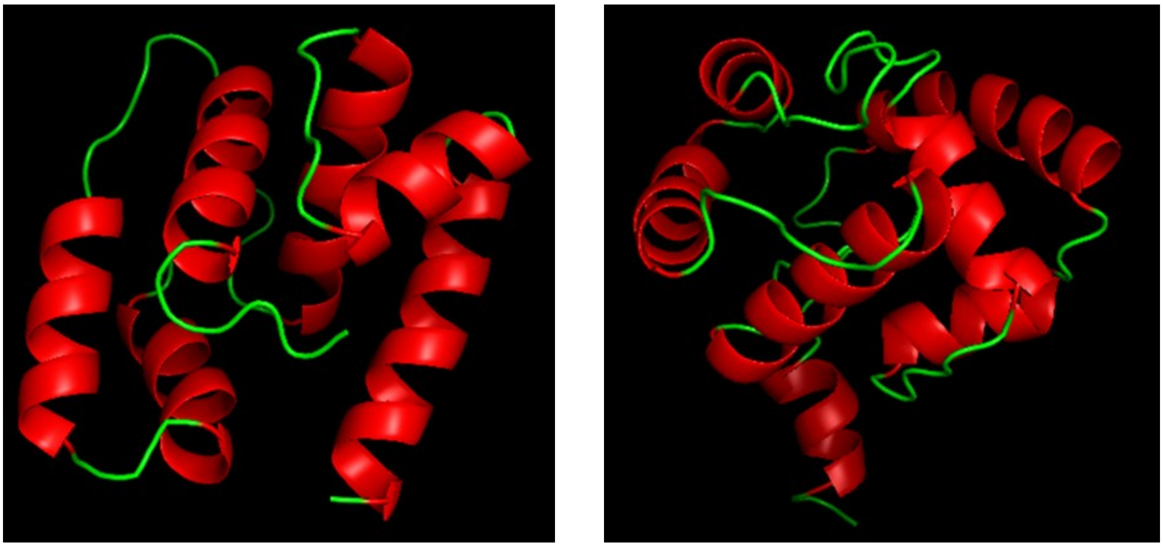

**Figure S1.** Cartoon representation Helix and Loop of the structure of RferOBP1766 (Left) and RferOBP23 (Right). The 3-D structures were model using a homology modelling tool 1-TASSER. The figures were rendered by using Pymol.

**Table S1.** The CASTp server was used to identify potential binding pockets in RrefOBP1768 and RrefOBP23 3-D structures. Amino acids involved in the main binding pockets (Active sites) of both proteins are listed.

| **RferOBP1768** | **Amino acid nature** | **RferOBP23** | **Amino acid nature** |
|---|---|---|---|
| M1 | Hydrophobic | F12 | Hydrophobic |
| Q4 | Hydrophilic | L13 | Hydrophobic |
| R5 | Positive | P14 | Hydrophobic |
| R7 | Positive | Y15 | Hydrophobic |
| F8 | Hydrophobic | C18 | Hydrophilic |
| F11 | Hydrophobic | I19 | Hydrophobic |
| Q12 | Hydrophilic | R49 | Positive |
| V26 | Hydrophobic | L88 | Hydrophobic |
| A20 | Hydrophobic | P89 | Hydrophobic |
| F30 | Hydrophobic | Y92 | Hydrophobic |
| G32 | Hydrophobic | T95 | Hydrophobic |
| L34 | Hydrophobic | N126 | Hydrophilic |
| H43 | Positive | P127 | Hydrophobic |
| L44 | Hydrophobic | A128 | Hydrophobic |
| V47 | Hydrophobic | H129 | Positive |
| G48 | Hydrophobic | Y130 | Hydrophobic |
| K50 | Positive | F131 | Hydrophobic |
| G51 | Hydrophobic | | |
| V53 | Hydrophobic | | |
| V64 | Hydrophobic | | |
| M65 | Hydrophobic | | |
| G68 | Hydrophobic | | |
| I69 | Hydrophobic | | |
| F72 | Hydrophobic | | |
| V73 | Hydrophobic | | |
| M82 | Hydrophobic | | |
| M101 | Hydrophobic | | |
| M104 | Hydrophobic | | |
| F105 | Hydrophobic | | |
| H108 | Positive | | |
| F109 | Hydrophobic | | |
| G110 | Hydrophobic | | |
| A111 | Hydrophobic | | |
| **Total** | 33/111 | Total | 17/122 |
| Positive | 5 = 5/33 = 15.1% | Positive | 2 = 2/17 = 11.8 |
| Hydrophilic | 2 = 2/33= 6.1 % | Hydrophilic | 2 = 2/17 = 11.8 |
| Negative | 0 | Negative | 0 |
| Hydrophobic | 26 = 26/33 = 78.8% | Hydrophobic | 13 = 13/17 = 76.4% |

**Table S2.** Potential Stable mutants around the binding pockets (active sites) of RferOBP23 and RferOBP1768, generated using PoPMuSiC Program.

| **RferOBP1768 Mutant** | **ΔΔG (Kcal/mol)** | **Original amino acid** | **New amino acid** | **Original amino acid-HI1** | **New amino acid-HI1** |
|---|---|---|---|---|---|
| M1L | -0.02 | Hydrophobic | Hydrophobic | 1.8 | 3.8 |
| Q4I | -0.33 | Hydrophilic | Hydrophobic | -3.5 | 4.5 |
| Q4L | -0.32 | Hydrophilic | Hydrophobic | -3.5 | 3.8 |
| Q4V | -0.07 | Hydrophilic | Hydrophobic | -3.5 | 4.2 |
| Q4M | -0.07 | Hydrophilic | Hydrophobic | -3.5 | 1.8 |
| R5L | -0.18 | Positive | Hydrophobic | -4.5 | 3.8 |
| R7I | -0.1 | Positive | Hydrophobic | -4.5 | 4.5 |
| R7L | -0.08 | Positive | Hydrophobic | -4.5 | 3.8 |
| Q12I | -0.55 | Hydrophilic | Hydrophobic | -3.5 | 4.5 |
| Q12F | -0.5 | Hydrophilic | Hydrophobic | -3.5 | 2.8 |
| Q12L | -0.48 | Hydrophilic | Hydrophobic | -3.5 | 3.8 |
| Q12M | -0.42 | Hydrophilic | Hydrophobic | -3.5 | 1.8 |
| Q12V | -0.3 | Hydrophilic | Hydrophobic | -3.5 | 4.2 |
| Q12W | -0.14 | Hydrophilic | Hydrophobic | -3.5 | -0.9 |
| Q12Y | -0.08 | Hydrophilic | Hydrophobic | -3.5 | -1.3 |
| H43Y | -0.16 | Positive | Hydrophobic | -3.2 | -1.3 |
| H43M | -0.12 | Positive | Hydrophobic | -3.2 | 1.8 |
| G48L | -0.38 | Hydrophobic | Hydrophobic | -0.4 | 3.8 |
| G48M | -0.19 | Hydrophobic | Hydrophobic | -0.4 | 1.8 |
| G48A | -0.17 | Hydrophobic | Hydrophobic | -0.4 | 1.9 |
| G48W | -0.05 | Hydrophobic | Hydrophobic | -0.4 | -0.9 |
| **Total mutants = 21** | | | | | |
| **RferOBP23 Mutant** | **DDG (Kcal/mol)** | **Original amino acid** | **New amino acid** | **Original amino acid -HI1** | **New amino acid-HI1** |
| R49L | -0.29 | Positive | Hydrophobic | -4.5 | 3.8 |
| T95V | -0.02 | Hydrophobic | Hydrophobic | -0.7 | 4.2 |

Total mutants = 2

HI = Hydrophobicity Index

1= https://doi.org/10.1016/0022-2836(82)90515-0

**Table S3.** Binding Pocket Features – RrefOBP1768 and RferOBP23.

| **Main binding pocket (Active site)** | **RferOBP1768** | **RrefOBP23** |
|---|---|---|
| Area (Solvent accessible surface - AS) Å$^2$ | 401.5 | 89.303 |
| Area (Molecular surface - MS) Å$^2$ | 850.066 | 292.452 |
| Volume (Solvent accessible surface - AS) Å$^3$ | 229.81 | 28.175 |
| Volume (Molecular surface - MS) Å$^3$ | 1046.368 | 284.821 |
| Pocket length (Å) | 381.709 | 132.595 |

| | | |
|---|---|---|
| **Number of mouth openings** | **1** | **1** |
| Mouth area (Solvent accessible surface - AS) Å$^2$ | 14.534 | 6.429 |
| Mouth area (Molecular surface - MS) Å$^2$ | 45.79 | 29.84 |
| Mouth Length (Solvent accessible surface - AS) Å | 18.615 | 12.322 |
| Mouth Length (Molecular surface - MS) Å | 27.41 | 21.12 |
| **Number of binding pocket residues/whole protein** | **33/111, (29.7%)** | **17/122, (13.9%)** |
| Binding pocket residues - Hydrophobic | 26(78.8%) | 13(76.4%) |
| Binding pocket residues - Hydrophilic | 2(6.1%) | 2(11.8%) |
| Binding pocket residues - Positive | 5(15.1%) | 2(11.8%) |
| Binding pocket residues - Negative | 0 | 0 |
| **Number of residues that gave stable mutants (PoPMuSiC)** | **7/33, (21.2%)** | **2/17, (11.8%)** |
| Residues that gave stable mutants - Hydrophobic | 2 (28.6%) | 1(50%) |
| Residues that gave stable mutants - Hydrophilic | 2(28.6%) | 0 |
| Residues that gave stable mutants - Positive | 3 (42.8%) | 1(50%) |
| Residues that gave stable mutants - Negative | 0 | 0 |
| **Total number of stable mutants** | **21** | **2** |
| Stable mutants - when a new residue is Hydrophobic | 21 (100%) | 2(100%) |
| Stable mutants - when a new residue is Hydrophilic | 0 | 0 |
| Stable mutants - when a new residue is Positive | 0 | 0 |
| Stable mutants - when a new residue is Negative | 0 | 0 |
| **Total number of mutant proteins with higher binding affinities than WT after docking screening with drug ligands** | **6** | **1** |
| Mutant proteins with higher binding affinities than WT after docking screening - Hydrophobic | 6(100%) | 1 (100%) |
| Mutant proteins with higher binding affinities than WT after docking screening - Hydrophilic | 0 | 0 |
| Mutant proteins with higher binding affinities than WT after docking screening - Positive | 0 | 0 |
| Mutant proteins with higher binding affinities than WT after docking screening - Negative | 0 | 0 |
| **Å = Angstrom** | | |

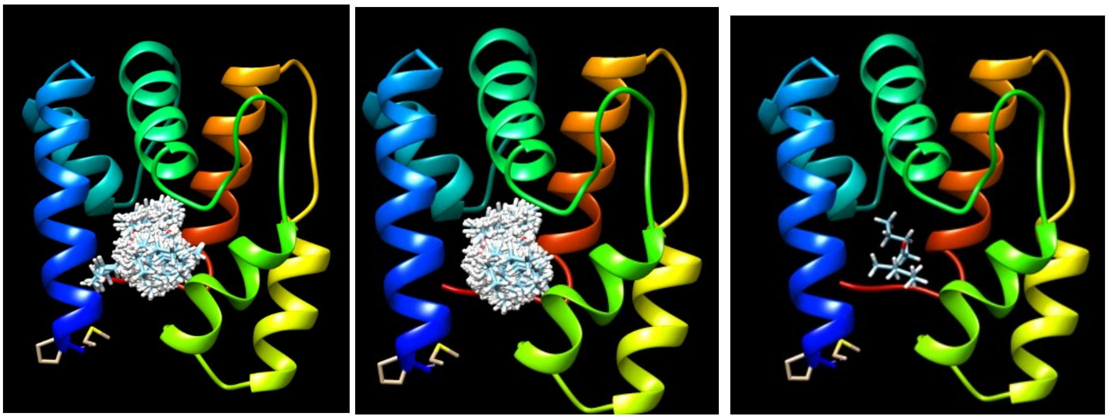

**(a)** **(b)** **(c)**

**Figure S2.** Visualisation of the docking outcomes using UCF Chimera. The docking outcome for RrefOBP1768 with a ferrugineone molecule is presented as an example. (a) all predicted clusters; (b) binding pocket clusters, (c) the most energetic cluster rank (binding mode) at the binding pocket. For docking experiments, target drug molecules in addition to the fluorescence probe 1-NPN were investigated. The latter is used in displacement binding experiments to determine the affinities of binding of target ligands. The docking tool was the Swissdock server (Swiss Institute of Bioinformatics (http://swissdock.vital-it.ch/docking) which operates using "EADock DSS software". Each of the potential stable mutants for RrefOBP1768 and RrefOBP23 **(Table S2)** plus the WTs was docked with each of the target ligands (**Supplementary Appendix 1)**. During the docking procedure, the software generates many binding modes simultaneously and their energies are estimated. The binding modes with the most favourable energies were evaluated and clustered. For each docking outcome up to 56 clusters were generated, each with several rankings, a cluster is a binding mode and within it there are several ranks, each rank has slightly different energy value compared to another rank, some of these ranks are just repetitions, a much bigger energy difference occurs between one binding mode to the other. The data were visualised using the plug in "UCF Chimera" and as reported in[15]. Initially all non-binding pocket clusters were eliminated. Then the most energetic binding pocket cluster was selected and refined by eliminating any repeated ranks and (for each ligand against each protein variant) this value was then recorded in **Supplementary Appendix 1**.

**Table S4.** Results of docking screening: This table presents potential stable mutants of RferOBP1768 and RferOBP23 (Table S2) that exhibit increased binding affinities towards target analytes compared to wild-type proteins. The underlined variants were cloned and expressed in the current study. For docking experiments, target ligands molecules in addition to the fluorescence probe 1-NPN were investigated (Cali et al., 2020). The latter is used in displacement binding experiments to determine the affinities of binding of target ligands. The actual docking was done using the Swissdock server (Swiss Institute of Bioinformatics (http://swissdock.vital-it.ch/docking) which operates using "EADock DSS software". Each of the potential stable mutants for RrefOBP1768 and RrefOBP23 (Table S2) plus the WTs was docked with each of the target ligands (**Supplementary Appendix 1)**. During the docking procedure, the software generates many binding modes simultaneously and their energies are estimated. The binding modes with the most favourable energies were evaluated and clustered. For each docking outcome up to 56 clusters were generated, each with several rankings, a cluster is a binding mode and within it there are several ranks, each rank has slightly different energy value compared to another rank, some of those ranks are just repetitions, a much bigger energy differences occurs between one binding mode to the other. The data were visualised using the plug in "UCF Chimera". Initially all non- binding pocket clusters were eliminated. Then the most energetic binding pocket cluster was selected and refined by eliminating any repeated ranks **(Figure S2)**. Binding energy values were used to define mutant variants with potentially stronger binding affinity towards a given ligand as the lower the binding energy the stronger the binding and vice versa. The mean value for the binding energy for the drugs for each protein variant was calculated, and the difference between the mean value of a given mutant and the WT mean value was recorded. Any difference value that was < 0 kcal/mol was used as an indicator of potentially stronger binding for that particular mutant than a WT (**Supplementary Appendix 1)** across all of the ligands tested[15].

| RferOBP23 variant | RferOBP1768 variant |
|---|---|
| R49L | Q4M<br>Q12I<br>Q12L<br>Q12M<br>Q12V<br>Q12W |

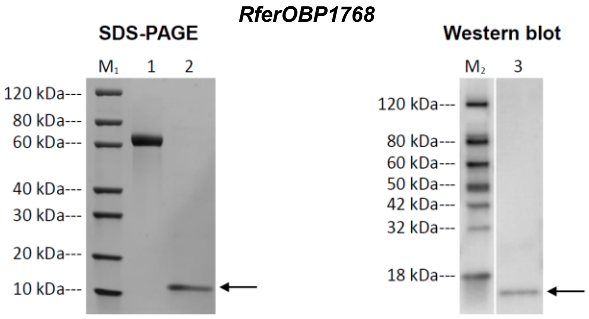


**Figure S3a.** SDS PAGE and WB analysis of RferOBP1768 peptide 22-132 AA (shown below). Lane M1: Protein Marker; Lane M2: Protein Marker; Lane 1: BSA (2.00 μg) Lane 2: RferOBP1768 23-132 AA; Lane 3: RferOBP1768 23-132 AA; MW: 13544.4. Primary antibody: Mouse-anti-His mAb. Arrows indicate the protein band.

```
                    10        20        30        40        50        60        70        80        90       100
            ....|....|....|....|....|....|....|....|....|....|....|....|....|....|....|....|....|....|....|....|
RferOBP1768 MCRFTAILLISLACGLIYGGMTPEQRTRFFNFQNECMQETGATDEMVLKAFAGELTDSPVFKDHLVCVGMKGGVIDEQGNFHKDVMKKGIMLFVDDEGKV

                   110       120       130
            ....|....|....|....|....|....|..
RferOBP1768 DAMLDKCYTHYDTQQDTAFNMMKCMFKEHFGA
```

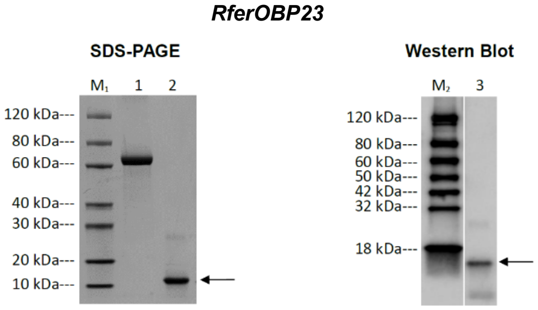


**Figure S3b.** SDS PAGE and WB analysis of RferOBP23 peptide 122 AA (shown below). Lane M1: Protein Marker; Lane M2: Protein Marker; Lane 1: BSA (2.00 μg) Lane 2: RferOBP23 122 AA (signal peptide removed); Lane 3: RferOBP23 122 AA. MW 13883.5. Primary antibody: Mouse-anti-His mAb. Arrows indicate the protein band.

```
                  10        20        30        40        50        60        70        80        90       100
          ....|....|....|....|....|....|....|....|....|....|....|....|....|....|....|....|....|....|....|....|
RferOBP23 MFKTLPIVLALFLPYISCISDEMKELAAQLHNACVAETGATEDAITNARAGTFADDDNFKCYFKCLFDQMAIMDDEGIIDVEAMIAVLPDEYQDTLPPVI

                 110       120       130
          ....|....|....|....|....|....|...
RferOBP23 RKCDTKKGANPCENAWLTHKCYYQENPAHYFLI
```

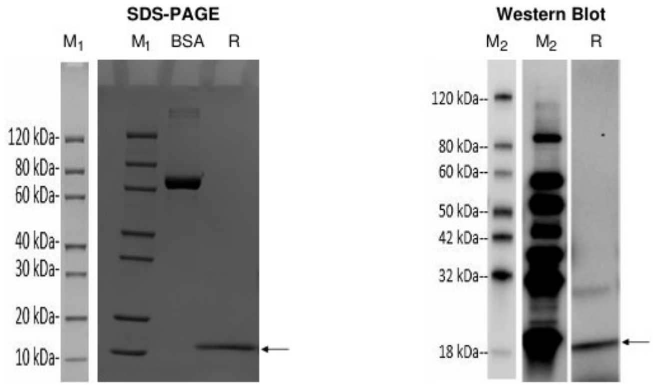


**Figuer S3c.** SDS PAGE and WB analysis of RferOBP23_R49L peptide 122 AA (shown below). Lane M1: Protein Marker; Lane M2: Protein Marker; Lane R: *RferOBP23_R49L* (Reducing condition). MW 13838.2.

MHHHHHHISDEMKELAAQLHNACVAETGATEDAITNALAGTFADDDNFKCYFKCLFDQMAIMDDEGII DVEAMIAVLPDEYQDTLPPVIR KCDTKKGANPCENAWLTHKCYYQENPAHYFLI . Primary antibody: Mouse-anti-His mAb.

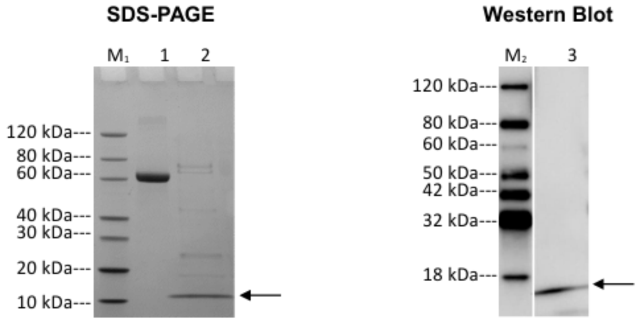


**Figure S3d.** SDS-PAGE and Western blot analysis of RferOBP1768_Q12V peptide 117 AA (shown below). Lane M1: Protein Marker, Lane M2: Protein Marker, Lane 1: BSA (2.00 μg), Lane 2: RferOBP1768_Q12V (Reducing condition), Lane 3: RferOBP1768_Q12V (Reducing condition). MW: MW=13513.0. Primary antibody: Mouse-anti-His mAb. Arrows indicate the protein band.

MHHHHHHPEQRTRFFNFVNECMQETGATDEMVLKAFAGELTDSPVFKDHLVCVGMKGGVI DEQGNFHKDVMKKGIMLFVDDEGKV DAMLDKCYTHYDTQQDTAFNMMKCMFKEHFGA

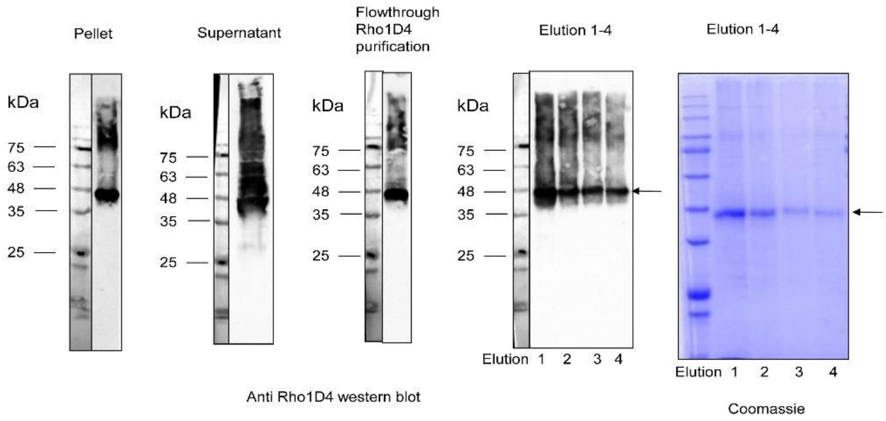


**Figure S4**. RferOR1 800µL upscale expression in the presence of 0.4 % Brij58; Rho1D4 purification, analysis on western blot and Coomassie blue stained SDS gel (elution fractions). Arrow indicates RferOR1 protein.

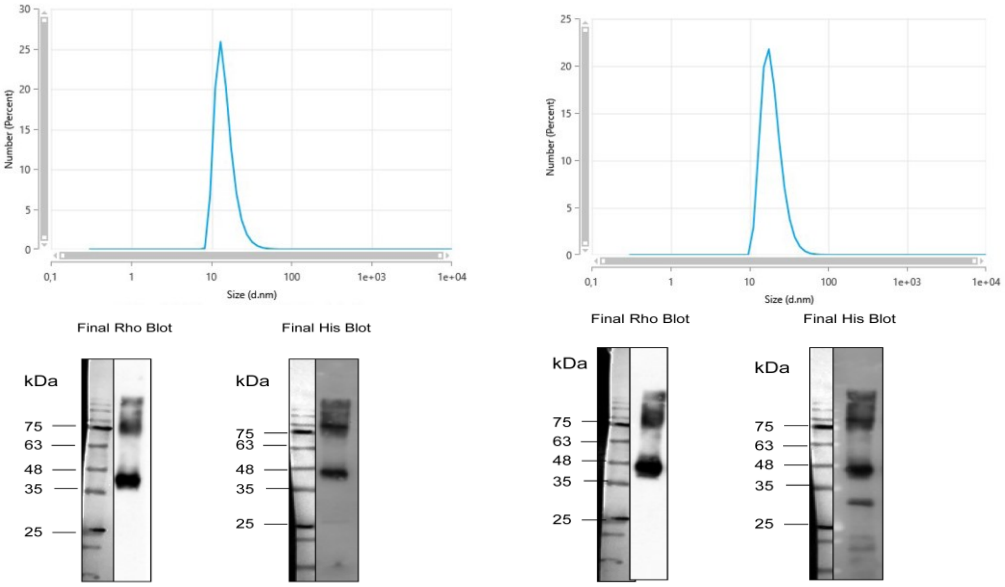


**Figure S5**. RferOR1 Upscale Expression and Integration Analysis. RferOR1 was expressed in a 2 mL upscale reaction with the presence of 0.4% Brij58. Following expression, the protein was integrated into two distinct membrane scaffold protein (MSP) nanodisc systems: MSP1D1 DMPC (left) (Mean by number (nm): 15,07) and MSP1E3D1 DMPC (right) (Mean by number (nm): 19,86). The quality and characteristics of the resulting nanodiscs were assessed using dynamic light scattering (DLS) measurements. Arrow indicates RferOR1 protein.

**Table S5.** Affinity constants determined experimentally in solution using fluorescence probe competitive binding assays.

| | **Affinity $K_a$ ($\mu M^{-1}$)** | | | |
|---|---|---|---|---|
| **Analyte** | **RferOBP1768** | **RferOBP1768_Q12V** | **RferOBP23** | **RferOBP23_R49L** |
| **1-NPN (Fluorescence probe)** | 3.39 ±2.0 | 0.56 ±0.02 | 2.27 ± 0.67 | 0.67 ± 0.12 |
| Ferrugineol | 1.36±0.54 | 0.10 + 0.01 | 0.58±0.19 | 0.05 + 0.01 |
| Ferrugineone | 1.73±0.17 | 0.06 + 0.00 | 3.08±1.06 | 0.05 + 0.01 |
| Isopropyl acetate | 0.52 + 0.08 | 0.12 + 0.01 | 0.09 + 0.01 | 0.05 + 0.00 |

| | | | | |
|---|---|---|---|---|
| α−farnesene | 2.70 + 0.37 | 0.71 + 0.08 | 0.33 + 0.02 | 0.19 + 0.02 |
| Limonene | 0.46 + 0.03 | 0.05 + 0.01 | 0.11 + 0.01 | 0.05 + 0.01 |
| α−Pinene | 0.38 + 0.03 | 0.13 + 0.03 | 0.12 + 0.01 | 0.08 + 0.01 |
| Ethyl acetate | 0.38 ±0.002 | 0.13 + 0.04 | 0.08 ±0.02 | 0.07 + 0.02 |
| Geraniol | 0.26 + 0.01 | 0.09 + 0.01 | 0.14 + 0.01 | 0.07 + 0.01 |
| β−Pinene | 0.27 + 0.01 | 0.10 + 0.02 | 0.08 + 0.01 | 0.06 + 0.00 |
| Linalool | 0.14 + 0.01 | 0.21 + 0.05 | 0.10 + 0.02 | 0.05 + 0.00 |
| Carvacrol | 0.18 + 0.03 | 0.19 + 0.01 | 0.09 + 0.02 | 0.09 + 0.00 |
| p-cymene | 0.19 + 0.07 | 0.16 + 0.03 | 0.16 + 0.01 | 0.06 + 0.02 |
| Furfuryl alcohol | 0.16 + 0.02 | 0.15 + 0.03 | 0.10 + 0.02 | 0.05 + 0.01 |
| Acetaldehyde | 0.33 + 0.06 | 0.19 + 0.04 | 0.13 + 0.02 | 0.06 + 0.01 |
| Octanal | 0.23 + 0.03 | 0.11 + 0.03 | 0.05 + 0.01 | 0.02 + 0.00 |
| Hexene-1-ol | 0.28 + 0.02 | 0.21 + 0.03 | 0.12 + 0.01 | 0.09 + 0.01 |
| Naphthalene | 0.19 + 0.01 | 0.12 + 0.03 | 0.09 + 0.01 | 0.09 + 0.01 |
| δ−valerolactone | 0.33 + 0.04 | 0.16 + 0.06 | 0.08 + 0.02 | 0.11 + 0.01 |
| 5-Methyl furfural | 0.31 + 0.06 | 0.15 + 0.02 | 0.11 + 0.01 | 0.09 + 0.01 |
| Camphor | 0.13 + 0.01 | 0.20 + 0.01 | 0.10 + 0.02 | 0.08 + 0.00 |
| Thymol | 0.12 + 0.01 | 0.15 + 0.01 | 0.10 + 0.01 | 0.09 + 0.01 |
| Menthol | 0.20 + 0.07 | 0.18 + 0.03 | 0.08 + 0.01 | 0.06 + 0.01 |
| γ-undecalactone | 0.46 + 0.02 | 0.17 + 0.03 | 0.08 + 0.02 | 0.11 + 0.01 |

**Figure S6**. Sensor Array preparation and test. OBPs and OR1 from RPW are used to functionalize a set of 20 MHz Quartz Microbalances. Six sensors, functionalized with wild-type and mutant OBPs and ORs, were placed in a sensor cell and connected to electronic circuits for resonant frequency measurement and acquisition. Sensors were kept under a constant flow of synthetic air, during which pulses of vapor were applied from a set of test compounds.

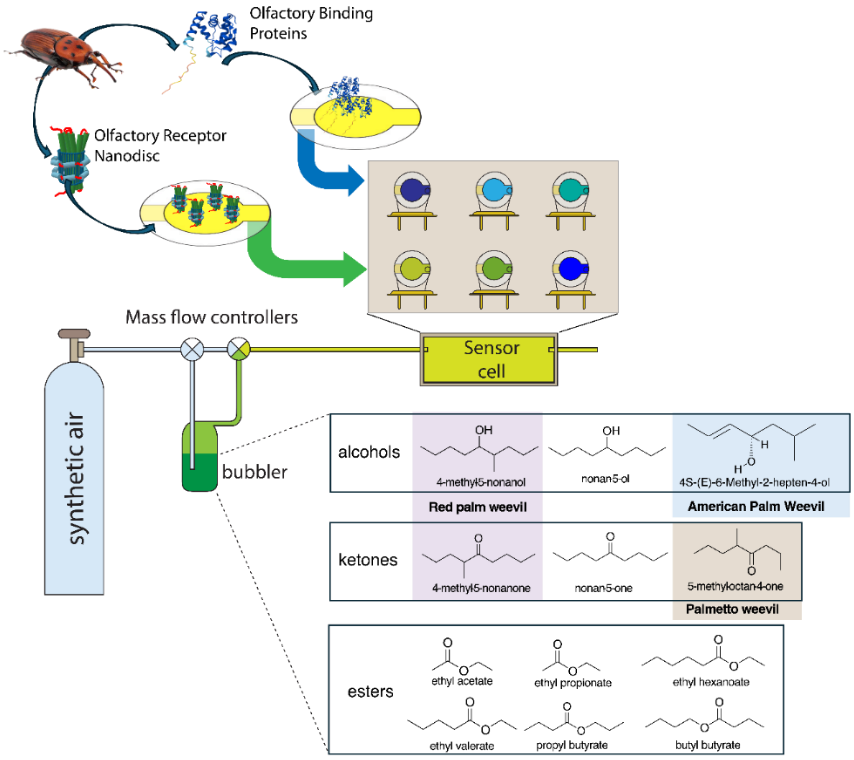


**Table S6.** Binding energies from the docking of the RWP OBPs used in the biosensor array development with the compounds in Table 2 OR Table S5. The lower the binding energy the higher the affinity and *vice versa*.

| | **Energy [E(Kcal/mol)]** | | | | |
|---|---|---|---|---|---|
| **Analyte** | **RferWTOBP1768** | **RferOBP1768_Q12V** | **RferWTOBP23** | **RferOBP23_R49L** | **ReferOR1** |
| ethyl hexanoate | -2.33 | -5.21 | 1.08 | 2.34 | -4.36 |
| ethyl valerate | -30.69 | -30.1 | -27.14 | -26.98 | -29.46 |
| butyl butyrate | -34.41 | -34.8 | -27.38 | -30.08 | -34.44 |
| propyl butyrate | -30.77 | -31.29 | -25.03 | -26.18 | -29.77 |
| nonan-5-one | -35.29 | -36.83 | -32.22 | -32.53 | -30.01 |
| nonan-5-ol | -34.22 | -35.46 | -27.85 | -28.46 | -35.93 |
| 5-methyloctan-4-one | -32.19 | -33.67 | -26.28 | -27.63 | -32.26 |
| ethyl butyrate | -27.21 | -27.6 | -20.33 | -22.66 | -22.24 |
| ethyl propionate | -21.62 | -21.62 | -17.21 | -20.35 | -18.8 |
| 2(E)-6-methyl-2-hepten-4-ol | -26.34 | -26.46 | -23.42 | -23.99 | -24.45 |
| Ferrugineol | -33.99 | -32.61 | -27.96 | -26.66 | -31.74 |
| Ferrugineone | -36.63 | -33.17 | -30.06 | -30.14 | -28.9 |
| Ethyl acetate | -18.68 | -18.86 | -15.15 | -16.37 | -15.33 |
